# A Spatiomolecular Map of the Striatum

**DOI:** 10.1101/613596

**Authors:** Antje Märtin, Daniela Calvigioni, Ourania Tzortzi, Janos Fuzik, Emil Wärnberg, Konstantinos Meletis

## Abstract

The striatum is organized into two major outputs formed by striatal projection neuron (SPN) subtypes with distinct molecular identities. In addition, the histochemical division into patch and matrix compartments represents an additional spatial organization, proposed to mirror a functional specialization in a motor-motivation dimension. To map the molecular diversity of SPNs in the context of the patch and matrix division, we genetically labeled mu-opioid receptor (Oprm1) expressing striatal neurons and performed single-nucleus RNA sequencing (snRNA-seq). This allowed us to establish new molecular definitions of the patch-matrix compartments, resulting in a molecular code for mapping patch SPNs at the cellular level. In addition, Oprm1 expression labeled exopatch SPNs, which we found to be molecularly distinct from both patch as well as neighboring matrix SPNs, thereby forming a separate molecular entity. At the cell-type level, we found an unexpected SPN diversity, leading to the identification of a new Col11a1+ striatonigral SPN type. At the tissue level, we found that mapping the spatial expression of a number of markers revealed new definitions of spatial domains in the striatum, which were conserved in the non-human primate brain. Interestingly, the spatial markers were cell-type independent and instead represented a spatial code that was found across all SPNs within a spatially restricted domain. This spatiomolecular map establishes a formal system for targeting and studying the striatal subregions and SPNs subtypes, beyond the classical striatonigral and striatopallidal division.

## INTRODUCTION

The organization of the striatal circuitry is based on the division of striatal projection neurons (SPNs) into two major pathways that have distinct neuroanatomical and molecular features^1^. In the established basal ganglia model^2,3^, the striatonigral SPNs (direct pathway) express dopamine D1 receptors (Drd1), whereas the striatopallidal SPNs (indirect pathway) express dopamine D2 receptors (Drd2); and these two pathways have opposing roles during behavior. However, this model has been challenged by recent findings revealing an unexpected complexity in the activity of the two major SPN subtypes during motor behavior, demonstrating that the activity patterns of SPNs do not simply map onto a simple D1-D2 segregation^4,5^. In light of this mismatch between the functional and neuroanatomical classification of the striatal circuitry, a more detailed understanding of the molecular and functional heterogeneity of SPNs can guide the refinement of the basal ganglia model.

In addition to the cell-centric definition of the striatal circuit organization, important neuroanatomical and functional differences have been described at the tissue level, which for example span across the mediolateral and dorsoventral axes of the striatum^6^. The striatal volume has in this context been functionally divided into sensorimotor, associative, and limbic regions; this functional division is in a broad sense represented by a corresponding spatial division into dorsolateral, dorsomedial, and ventral domains of the striatum^7,8^. A major challenge remains in terms of formulating precise spatial definitions of the striatal subregion borders, since the molecular classification to definitely map these borders has not been established.

Another level of striatal tissue classification has been based on the differential distribution of either acetylcholinesterase activity or of mu opioid receptor (MOR) levels, representing a histochemical differentiation of the striatum into subregions^9,10,11,12^. This histochemical division into MOR-labeled striatal patches (also known as striosomes) and the surrounding striatal matrix is likely of functional relevance since it overlaps with segregation of input-output patterns, for example discrete cortical inputs^13,14^, as well as the topography of striatal outputs^15,16^. The patch compartment receives distinct projections from the medial and orbital prefrontal cortex and has therefore been categorized as part of a limbic circuit, whereas the matrix compartment – constituting the largest part of the striatal volume – based on its connectivity with sensorimotor areas has been primarily linked to action selection and motor control^17,18^.

We aimed in this study to define the molecular identity of patch and matrix SPNs and use this information to map the molecular composition of SPN subtypes in dorsal striatum. The unbiased molecular classification of neuron subtypes based on their gene expression pattern is now possible through single-cell RNA sequencing methods (scRNA-seq)^19,20^. This has formed the basis for a systematic classification of neuron types, and large efforts to map the diversity of cell types and tissue organization^21^. Since a major neuroanatomical and histochemical division of the striatum has been based on detection of MOR expression, we developed a genetic labeling approach using recombinase-mediated labeling of MOR-expressing cells (Oprm1+) in a new transgenic mouse line (Oprm1-Cre). The genetic access allowed us to isolate Oprm1+ striatal neurons and map their identity using single-nucleus RNA sequencing (snRNA-seq) and furthermore compare their molecular profile to SPNs in the matrix compartment. We show that the patch-matrix division can be defined at the cellular level using new molecular markers, and that in addition to the major D1+ and D2+ SPNs there are additional SPN subtypes. We have identified a discrete marker profile to define exopatch SPNs and we describe a new Col11a1+ SPN subtype. In addition to the cell-type specific markers, we further describe the existence of specific spatial markers that allow us to define subregions of the striatum based on gene expression.

In summary, we present a spatiomolecular map of the striatum, based on snRNA-seq and spatial mapping of gene expression, that is the foundation for experimental structure-function dissection of the striatum.

## RESULTS

### Genetic Labeling and Mapping of Oprm1+ SPNs

To identify striatal neurons belonging to the patch compartment at the cellular level, we genetically labeled the Oprm1+ striatal neurons by generating an Oprm1-2A-Cre knock-in mouse line (Oprm1-Cre). This Oprm1-Cre mouse line allowed us to genetically target and identify Oprm1+ striatal neurons using Cre-mediated recombination. To visualize Oprm1+ striatal neurons, we crossed the Oprm1-Cre mouse with a Cre-dependent reporter mouse line (Ai14 reporter mouse line; R26R-tdTomato^22^) to generate mice with tdTomato expression in Oprm1+ striatal neurons (Oprm1:tdTomato). The Oprm1:tdTomato mice allowed us to visualize and map the Oprm1+ striatal neurons (Fig. 1A). We first used immunostaining against MOR to visualize the striatal patches and map the distribution of tdTomato+ cells in patches versus in matrix. We found that the density of tdTomato+ cells was higher in MOR-dense patches compared to the surrounding tissue in four different anteroposterior coordinates (Fig. 1B).

**Figure 1.**
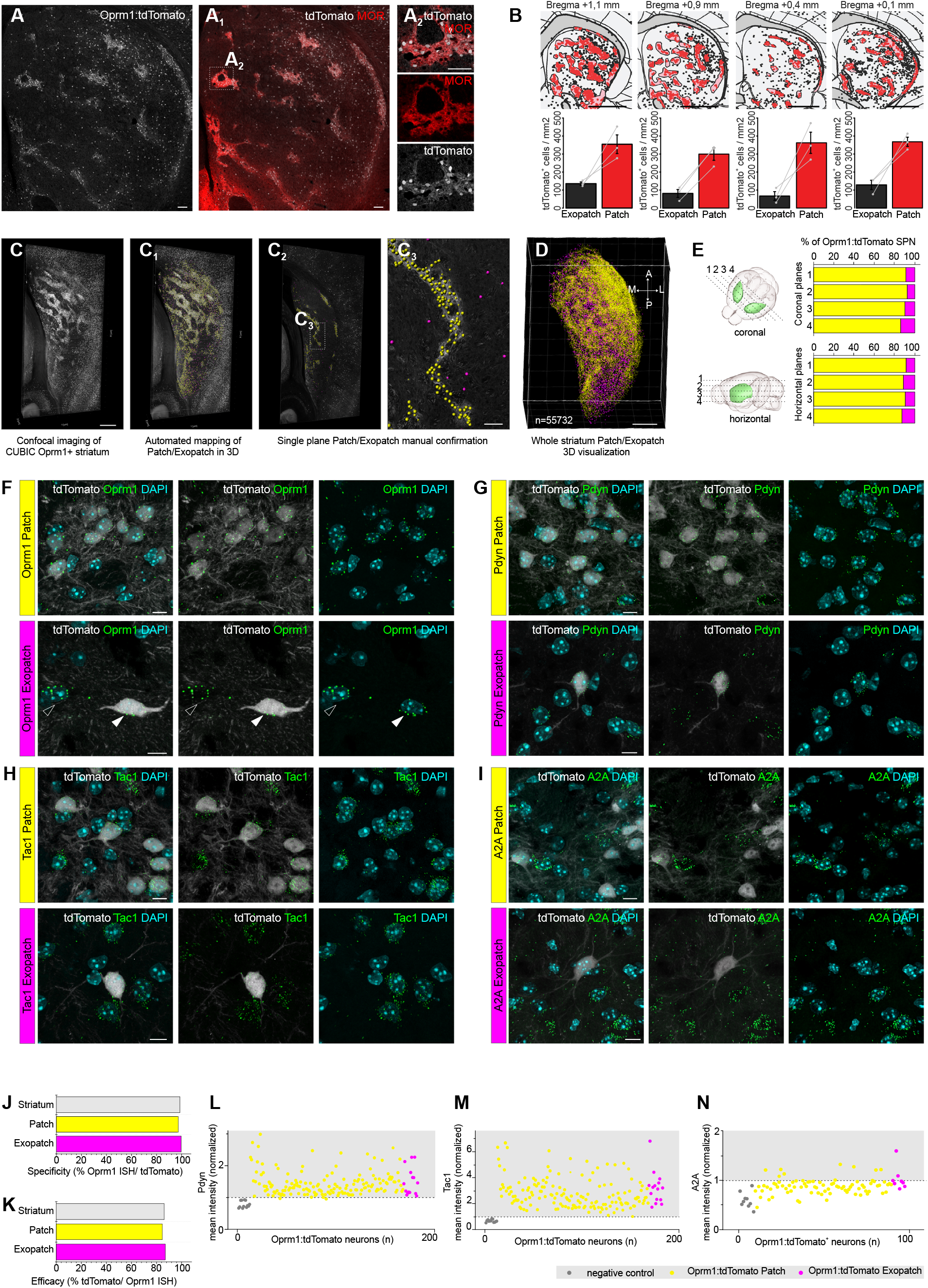
Mapping of Patch SPNs and Exopatch SPNs in Oprm1-Cre:tdTomato mice. **(A)** Oprm1-Cre dependent tdTomato expression (*white*) labels striatal neurons clustered in patches, defined by high mu opioid receptor (MOR) immunostaining (*red*) **(A1-A2)**, as well as lower number of sparse neurons within the matrix, defined as lower MOR immunostaining. (B) Digital reconstruction of patch (pink background) and matrix (white background) striatal areas defined by respectively high and low MOR immunolabeling. Segmented patch Oprm1-Cre:tdTomato SPNs are shown in red and the exopatch SPNs in black (top row). Relative cell density (cells/mm^2^) in the patch versus the exopatch compartment in four striatal coordinates (bottom row, n=3, two sample t-test, p<0,01). **(C)** 3D visualization of Oprm1-Cre:tdTomato SPNs (*white*) in 750 μm thick CUBIC cleared horizontal section of the striatum. Oprm1-Cre:tdTomato SPNs were automatically detected and spatially classified as Patch SPNs (*yellow*) or Exopatch SPNs (*magenta*) within the 3D volume **(C1)**. Automated classification in the 3D volume was confirmed by an expert-based single plane validation **(C2-C3). (D)** Visualization of Patch SPNs and Exopatch SPNs in the whole striatum from four horizontal sections (750 μm thick CUBIC cleared sections were aligned, showing 55732 segmented neurons) **(C4). (E)** Quantification of the proportion of Patch SPNs (*yellow*) and Exopatch SPNs (*magenta*) in the whole striatum (*green in the schematic*) calculated in four 750 μm thick coronal sections or four 750 μm thick horizontal sections (illustrated as dashed lines in the schematic). **(F)** RNAscope ISH revealed high expression of Oprm1 (*green*) both in Oprm1:tdTomato-positive (*white*) Patch SPNs (*yellow*, upper row) and Exopatch SPNs (*magenta*, lower row, *full arrowhead*). Both Oprm1:tdTomato-positive (*white*) Patch SPNs and Exopatch SPNs expressed high level of Prodynorphin (**G**, *green*), Tac1 (**H**, *green*) but did not express Adora2A (**I**, *green*). **(J)** Specificity of Oprm1-Cre:tdTomato mouse line determined as proportion of Oprm1 (ISH)-positive cells and Oprm1-Cre:tdTomato positive neurons in Patch SPNs (*yellow*), Exopatch SPNs (*magenta*) and total Oprm1-Cre:tdTomato positive population. **(K)** Efficacy of Oprm1-Cre:tdTomato mouse line determined as proportion of tdTomato-positive neurons and Oprm1 (ISH) positive cells in Patch SPNs (*yellow*), Exopatch SPNs (*magenta*) and total Oprm1-Cre:tdTomato positive population. (**L-N**) Quantification of Prodynorphin, Tac1 and Adora2A RNA expression levels in Oprm1-Cre:tdTomato negative cells (negative controls, *gray*), Patch SPNs (*yellow*) and Exopatch SPNs (*magenta*). Data shows single cell normalized mean intensity. Threshold for positive expression (*dashed line*) was set for each probe to two standard deviations above the mean for the negative control. Scale bar: A 100μm, B 1 mm, C-C_2_ 500 μm, C_3_ 100 μm, D 1 mm, F-I 10 μm. N=animals number, n= cells number.

In addition to the tdTomato+ cells in patches, we detected tdTomato+ cells distributed in the matrix compartment, found throughout the striatum but primarily enriched in the dorsolateral domain. We adopted the nomenclature “exopatch” cells from a recent study^23^ to denote Oprm1:tdTomato+ cells that reside outside the densely MOR-labeled patch compartment. To visualize the organization of Oprm1+ cells in the striatal tissue, we applied a tissue clearing protocol in Oprm1:tdTomato mice (CUBIC^24^) and then mapped the labeled Oprm1+ striatal neurons and patches in 3D (Fig. 1C-D). The 3D visualization of Oprm1:tdTomato+ neurons allowed us to define patches using an unbiased classification, based solely on the spatial relationship between Oprm1:tdTomato+ cells (Video 1). We quantified the 3D relationship between Oprm1:tdTomato+ cells based on their distance to the five nearest Oprm1:tdTomato+ neighboring cells (Fig. 1E and Suppl. Fig 1-3). Based on this volume relationship, we determined that 88.5 % of Oprm1:tdTomato+ cells can be classified as belonging to the patch compartment.

To characterize the specificity and identity of the Oprm1:tdTomato cell labeling, we performed in situ hybridization (ISH) to quantify the presence of Oprm1 mRNA in tdTomato+ cells (Fig. 5F-I). We found that most Oprm1:tdTomato+ striatal cells were positive for Oprm1 expression in both the patch (patch SPNs) as well as matrix (exopatch SPNs) compartment (Fig. 1F, J). Importantly, we were able to genetically capture most of the Oprm1-expressing cells using the Oprm1-Cre mouse line (Fig.1K). We could further determine that most Oprm1:tdTomato+ cells displayed characteristic marker expression of the D1-type SPN, including Pdyn and Tac1, determined by in situ hybridization (Fig. 1G-I, L-N). In summary, the tissue mapping and marker quantification establishes that Oprm1:tdTomato+ cells primarily represent the Oprm1+ patch SPNs, but also capture the Oprm1+ exopatch SPNs.

### Molecular Classification SPNs in Patch, Exopatch, and Matrix

In order to define the molecular diversity of SPN, and more specifically to identify cell-type specific markers of patch and exopatch SPNs, we use a single nucleus RNA-seq (snRNA-seq) method to map RNA expression in single SPNs. We isolated and sorted the nuclei of single SPNs, defined as GABAergic or Oprm1+, using a nuclear isolation protocol^25^ in combination with fluorescence activated cell sorting (FACS) from the dorsal striatum of adult Vgat-Cre:H2B-GFP or Oprm1-Cre:H2B-GFP mice. We used a Smart-seq2 protocol^26^ to define the RNA expression profile in single nuclei with high sensitivity, resulting after quality control in the detection of 3764 unique genes per nucleus on (average 187160 reads per nucleus) from 375 Vgat-Cre:H2B-GFP nuclei and 351 Oprm1-Cre:H2B-GFP nuclei (Suppl. Fig. 4).

We first visualized in the expression of general neuronal and striatal markers in t-SNE plots, for example the striatal marker Ppp1r1b (also known as DARPP-32, expressed in 99% of nuclei, Suppl. Fig. 5), and the expression of cell-type specific markers to define the identity of the major clusters, based on the division of SPNs into D1+ (Drd1) and D2+ (Drd2, Adora2a) populations (Fig. 2A-C, Suppl. Fig. 5). This classification allowed us to map the majority of sequenced striatal nuclei into either Drd1+ or Drd2+/Adora2+ clusters (94% of the nuclei), representing the classical D1-D2 molecular division of SPNs. In addition, we identified three small clusters with gene expression patterns that suggested striatal interneuron identity^27^ (cluster 27: Npy+, Nxph2+; cluster 28: Th+; cluster 29: Pvalb+, Pthlh+) (Suppl. Fig 5). We found that the majority of Oprm1-Cre:H2B-GFP nuclei expressed markers for D1+ SPNs (87% defined as D1+, 9% defined as D2+), whereas Vgat-Cre:H2B-GFP nuclei could be classified as either D1+ or D2+ (57% defined as D1+, 37% defined as D2+). The snRNA-seq data therefore support a cell-type classification model where Oprm1+ neurons belong primarily to the D1+ SPN subtype.

**Figure 2.**
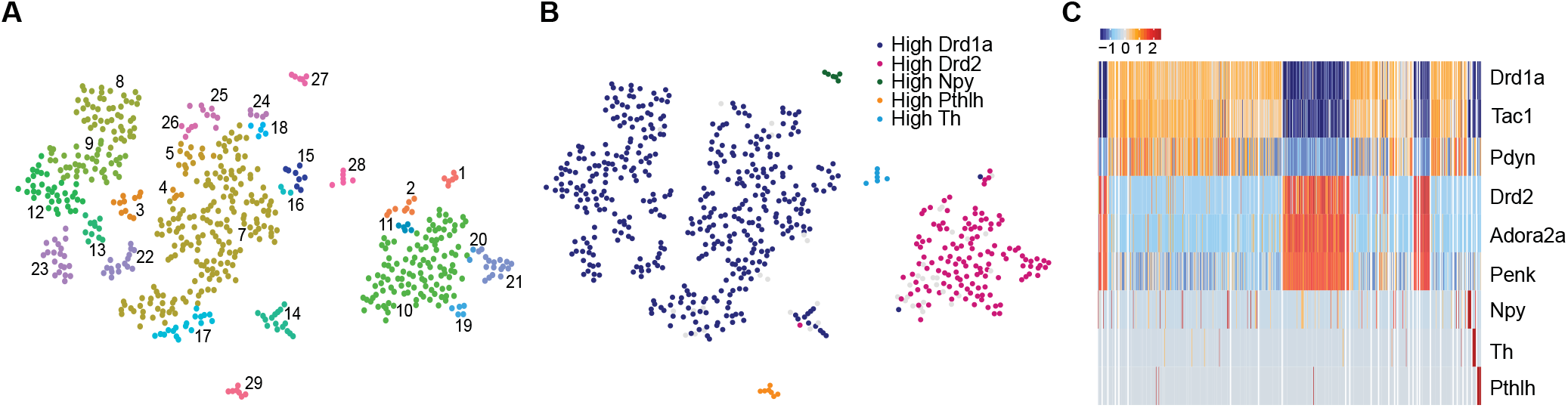
Mapping the molecular diversity of striatal neurons based on snRNA-seq. **(A)** Visualization of the molecular diversity of striatal neurons in a t-SNE plot showing the 29 clusters. Each dot represents an individual striatal neuron nucleus. **(B)** Mapping the identity of the major clusters based on established striatal molecular subtypes (D1+ SPN: Drd1a; D2+ SPN: Drd2; and three interneuron subtypes: Npy, Pthlh, Th). **(C)** Heatmap showing the expression of established markers for the classical SPNs and interneurons in each cluster for individual neuron nuclei (z-score of normalized log expression).

We further aimed to identify new molecular markers in our snRNA-seq data to define the striatal patch-matrix division, and for that reason we mapped the expression of enriched markers in candidate patch clusters (clusters 5, 7, 22). We found a number of markers preferentially labeling patch clusters (e.g. Sema5b, Mfge8, Kremen1, Pde1c) versus markers labeling matrix clusters (e.g. Id4, Sgk1, Epha4, Sv2b) (Fig. 3A-B, Suppl. Fig. 6-7), allowing us to potentially distinguish between patch and matrix striatal compartments by mapping gene expression with cellular resolution. We analyzed the expression pattern for these candidate patch-matrix markers in the adult striatum using the mouse in situ expression data in the Allen Brain Atlas^28^, and this confirmed a spatially segregated expression pattern that reflected the expected pattern of the striatal patch organization (Fig. 3C-D, Suppl. Fig 6). A recent study reported enriched expression of Asic4 in D2+ patch SPNs, and Necab1 expression in D1+ patch SPNs^30^. In our snRNA-seq data, Asic4 expression predominantly labeled D1+ SPNs in both patch and matrix, and Necab1 was enriched in the D2+ SPNs, although it was also expressed in D1+ SPNs and the Col11a1+ SPNs (Suppl. Fig 6).

**Figure 3.**
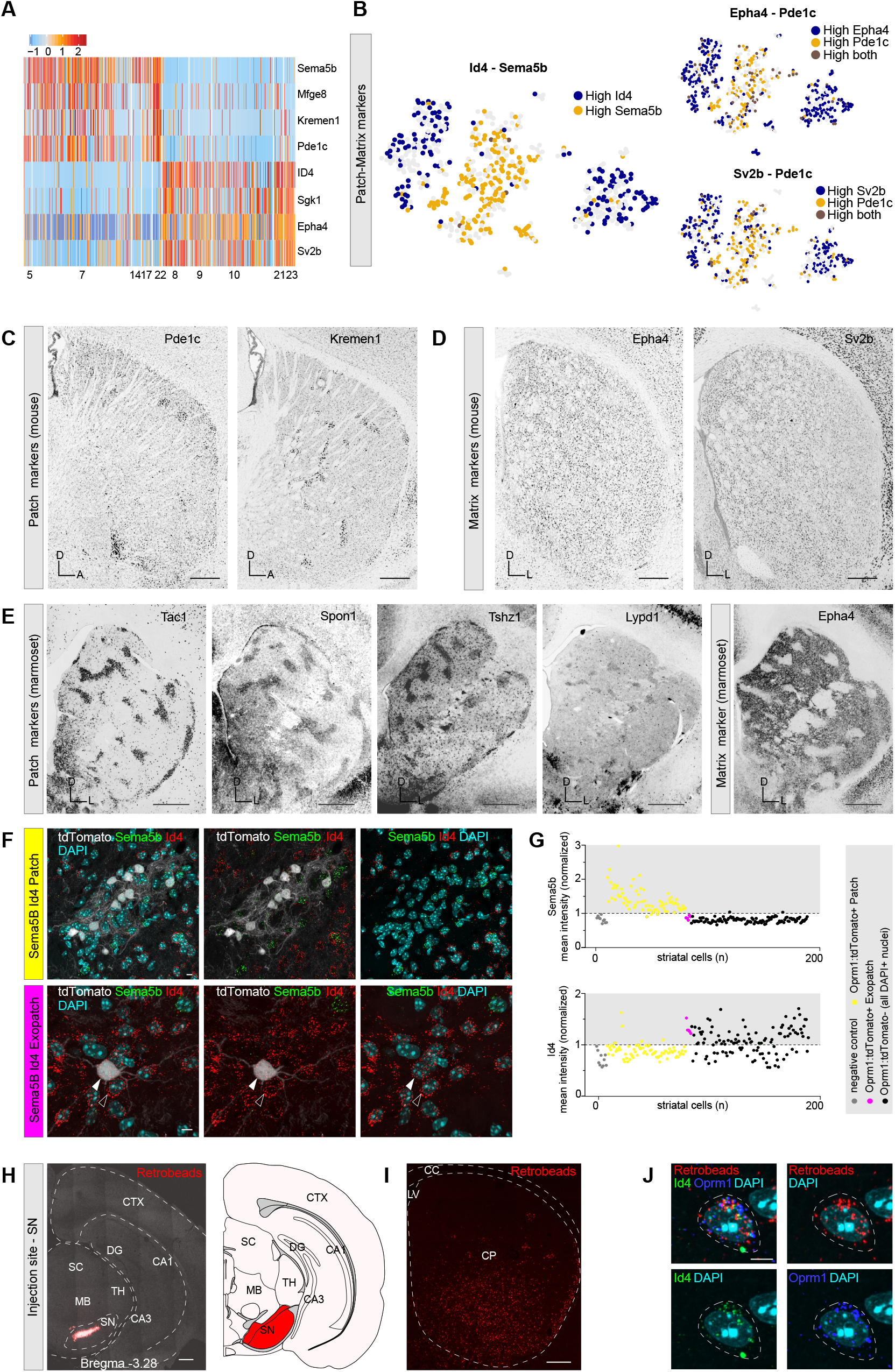
The molecular identity of Patch, Exopatch and Matrix SPNs. **(A)** Heatmap showing the classification of patch and matrix clusters based on new found markers (z-score of normalized log expression). Shown are five clusters identified as patch versus five clusters identified as matrix. **(B)** Distribution of expression of identified patch and matrix markers on t-SNE plot. Matrix markers are depicted in dark blue, patch markers in yellow. **(C)** Overview ISH images from Allen Brain Atlas show Pde1c and Kremen1 enriched expression in putative patches in the mouse brain. **(D)** Overview ISH images from Allen Brain Atlas show sparse distribution of striatal cells expressing Epha4 and Sv2b marking the putative matrix compartment in the mouse brain. **(E)** Overview ISH images from Marmoset Gene Atlas reveal Tac1, Spon1, Tshz1, Lypd1 enriched expression in putative patches and Epha4 high expression in putative matrix compartment in marmoset brain. **(F)** Sema5b (*green*) and Id4 (r*ed*) ISH on Oprm1-Cre:tdTomato mouse. Combining Sema5b and Id4 novel markers is sufficient to distinguish: Patch SPNs (*upper panels*) Oprm1-Cre:tdTomato^+^/Sema5b^+^/Id4^−^; Exopatch SPNs (*lower panels*) Oprm1-Cre:tdTomato^+^/Sema5b^−^/Id4^+^ (*full arrowhead*) and Matrix SPNs (*empty arrowhead*) Oprm1-Cre:tdTomato^−^/Sema5b^+^/Id4^−^. **(G)** Quantification of Sema5b and ID4 expression levels visualized as single cell normalized mean intensity in negative controls (*gray*), in Oprm1-Cre:tdTomato positive Patch SPNs (*yellow*), in Oprm1-Cre:tdTomato positive Exopatch SPNs (*magenta*) and Oprm1-Cre:tdTomato negative cells (*black*). Threshold for expression of each probe was set to the mean plus twice the standard deviation of the negative controls (*dashed line*). **(H)** Overview (*left*) and schematic (*right*) of Retrobeads injection site (*red*) in the substania nigra (SN). **(I)** Representative overview of SPNs (*red*) retrogradely labelled by Retrobeads injection to the SN. **(J)** High resolution example of long range projecting Exopatch SPNs retrogradely labelled after injection of retrobeads into substantia nigra (SN, *red*) co-expressing Oprm1 (ISH, *blue*) and Id4 (ISH, *green*). Abbreviations: (CA1) Cornu Ammonis subfield 1, (CA3) Cornu Ammonis subfield 3, (CC) corpus callosum, (CP) caudoputamen, (CTX) cortex, (DG) dentate gyrus, (MB) midbrain, (LV) lateral ventricle, (TH) thalamus, (SC) superior colliculus, (SN) substantia nigra. Scale bar: C-D 200 μm, E 1mm, F 10 μm, H-I 200 μm, J 10 μm. n= cells number.

To further determine whether the candidate patch-matrix markers displayed conserved expression patterns in non-human primates, we investigated the in situ expression in the corresponding dorsal striatum (i.e. the caudate and putamen) of the common marmoset (Callithrix jacchus) using the Marmoset Gene Atlas^29^. For the available markers (patch markers: Tac1, Spon1, Tshz1, Lypd1; matrix marker: Epha4), we found that they all displayed the predicted spatial expression pattern, labeling putative patch and matrix compartments in the marmoset caudate and putamen (Fig. 3E) To map in detail the expression pattern of two candidate patch-matrix markers (i.e. Sema5b, Id4), we mapped the in situ expression at cellular resolution in the dorsal striatum of Oprm1:tdTomato mice (Fig. 3F-G). We found that Sema5b expression labeled the majority of Oprm1:tdTomato+ patch cells, confirming the expression patter in our snRNA-seq data. In contrast, Oprm1:tdTomato-negative cells as well as exopatch Oprm1:tdTomato+ cells lacked Sema5b expression. In comparison, expression of Id4 labeled exopatch Oprm1:tdTomato+ and approximately half of Oprm1:tdTomato-negative cells (out of all cells identified by nuclear DAPI staining), whereas the Oprm1:tdTomato+ patch cells were negative for Id4 (Fig. 3G). We therefore conclude that Sema5b expression can molecularly define Oprm1+ striatal cells found in the patch compartment, and that Id4 is a marker for matrix striatal cells as well as for exopatch Oprm1+ striatal cells.

To further confirm that Id4 labels exopatch SPNs defined by their projection pattern, we performed labeling of the striatonigral projection through injections of retrograde-labeling beads (retrobeads) into the substantia nigra (SN) (Fig. 3H-I). This retrograde labeling allowed us to neuroanatomically define the Oprm1+ striatal cells as projection neurons, and we found that single Id4+ and Oprm1+ cells in dorsal striatum contained retrobeads (Fig. 3J). We therefore conclude that exopatch SPNs can be defined based on a unique molecular identity: co-expression of Id4 and Oprm1.

### Col11a1 Expression Defines a SPN subtype

In addition to the expected major D1+ and D2+ SPN populations, we also found other minor D1+ clusters in the snRNA-seq data. We examined the expression levels for well-established as well as novel markers to identify potentially novel SPN subtypes (Fig. 4A). We focused on cluster 14 (2.2% of the cells), which displayed expression of classical SPN markers (e.g. Drd1, Ppp1r1b), but was molecularly distinct from the larger Drd1+ and Adora2a+ clusters, and marked by distinct expression of primarily the markers Col11a1, Otof, Pcdh8, and Cacng5 (Fig. 4A-B). This cluster displayed marker expression similar to a striatal cluster in a recent study^30^, which classified these neurons as “eccentric” SPNs (eSPN), and defined them as a separate SPN population based on coexpression of Casz1, Otof, Cacng5, Pcdh8. The marker Otof was described as a marker for eSPNs^30^, and it labels Col11a1+ SPNs in our snRNA-seq data, although it also labels the cluster containing Npy+ striatal interneurons (Fig. 4B). In our snRNA-seq data, we found that cluster 14 was enriched for expression of Col11a1, Otof, Olfm3 (also found sparsely in other clusters), Pcdh8 (also found in Pvalb+ cluster), Adarb2, Sema5b, Cacng5, Fgfr1, Nxph4, Ralyl (Suppl. Fig. 8). In addition, nuclei in cluster 14 displayed on average high levels of Tac1 and Drd1 expression, and were negative for interneuron markers. This discrete gene expression profile suggested that the Col11a1+ cells represent a new SPN subtype.

**Figure 4.**
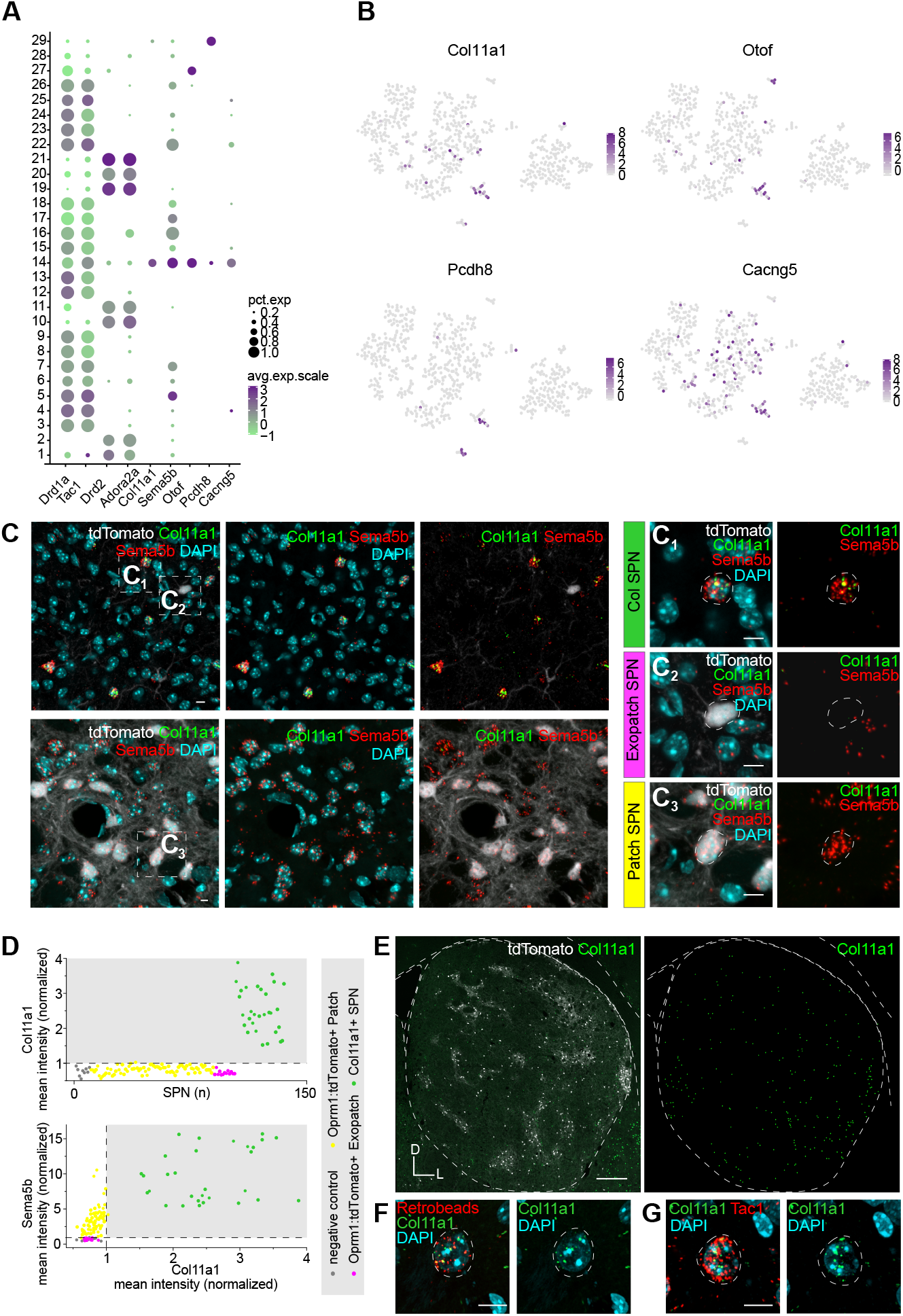
Col11a1 expression defines a SPN subtype with distinct molecular identity. **(A)** Dot-plot showing the expression of new identified cell type markers in cluster 14 compared to their expression in D1 and D2 SPNs. Dot size indicates fraction of cells in one cluster that express the gene. Dot color describes the z-score of the average expression value (of detected, non-zeros). **(B)** Distribution of expression of identified cell type markers on t-SNE plot (normalized log2-expression). **(C)** High resolution images reveal a sparse population of Col11a1 (arrowhead, upper panels) expressing SPNs. **(C_1_)** Col11a1 positive SPNs are identified as Col11a1^+^ (*green*)/ Sema5b^+^ (*red*)/ Oprm1-Cre:tdTomato^−^ (*white*). **(C_2_)** Col11a1 SPNs differ from Exopatch SPNs, identified as Oprm1-Cre:tdTomato^−^/ Col11a1^−^/Sema5b^−^, and from Patch SPNs **(C_3_)** identified as Oprm1-Cre:tdTomato^+^/ Col11a1^−^/ Sema5b^+^. **(D)** Col11a expression levels (*upper panel*) and Sema5b versus Col11a1 expression levels (*lower panel*) in negative controls (*gray*), in Patch SPNs (*yellow*), in Exopatch SPNs (*magenta*) and in Oprm1-Cre:tdTomato^−^/Col11a1^+^ SPNs (*green*). Threshold for expression of each probe was set to the mean plus twice the standard deviation of the negative controls (*dashed line*). **(E)** Representative overview (*left panel*) and segmented Col11a1 positive cells (*right panel*) reveal the sparse distribution of Col11a1 positive SPNs (ISH, *green*) outside Oprm1-Cre:tdTomato positive patches (*white*). **(F)** Col11a1 positive (ISH, *green*) long-range projecting SPNs retrogradely labelled after injection of retrobeads into substantia nigra (SN, *red*). **(G)** Tac1 expression (ISH, *red*) in Col11a1 positive SPNs (ISH, *green*). Scale bar: C-C_3_ 10 μm, E 200 μm, F-G 10 μm. n= cells number.

**Figure 5.**
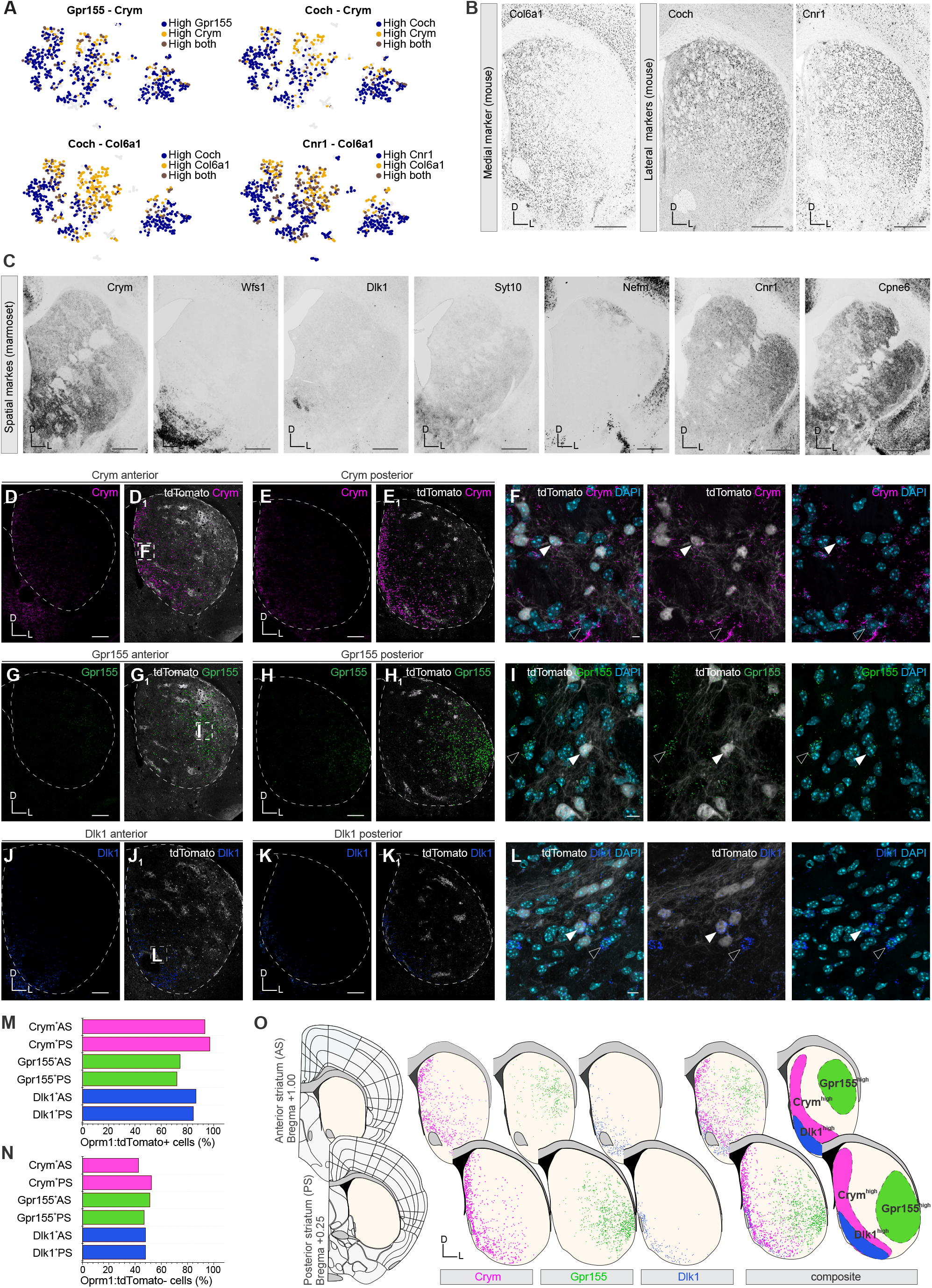
The striatum is defined by a spatiomolecular code. **(A)** Distribution of expression of identified spatial markers on t-SNE plot. More lateral markers are depicted in dark blue, more medial markers in yellow. **(B)** Overview ISH images from Allen Brain Atlas confirm high expression of Col6a1 in the medial striatum and high expression of the spatial markers Coch and Cnr1 in the lateral striatum of the mouse brain. **(C)** Overview ISH images from Marmoset Gene Atlas reveal conserved spatial distribution of Crym, Wfs1, Dlk1, Syt10, Nefm, Cnr1, Cpne6 in the marmoset caudate and putamen. **(D-E_1_)** Representative overviews of the anterior **(D-D_1_)** and the posterior **(E-E_1_)** striatum in Oprm1-Cre:tdTomato mice (*white*) probed by ISH for Crym (*magenta, ISH raw signal in left panels*). Segmented Crym^high^-expressing cells define a distinct area of the medial striatum (*magenta, segmented high-expressing SPNs in right panels*). **(F)** Within the Crym^high^-expressing area of the medial striatum Crym (*magenta*) is expressed by the Oprm1-Cre:tdTomato positive (*white*, full arrowhead) as well as Oprm1-Cre:tdTomato negative SPNs (empty arrowhead). **(G-H_1_)** Representative overview of the anterior **(G-G_1_)** and the posterior **(H-H_1_)** striatum of Oprm1-Cre:tdTomato mouse line (*white*) probed by ISH for Gpr155 (*green, ISH raw signal in left panels*). Distribution of the segmented Gpr155^high^-expressing cells define a distinct area of the lateral striatum (*green, segmented high-expressing SPNs in right panels*). **(I)** Within the Gpr155^high^-expressing area of the lateral striatum, the marker Gpr155 (*green*) is expressed by Oprm1-Cre:tdTomato positive (*white, full arrowhead*) as well as Oprm1-Cre:tdTomato negative SPNs (*empty arrowhead*). **(J-K_1_)** Representative overview of the anterior **(J-J_1_)** and posterior **(K-K_1_)** striatum from the Oprm1-Cre:tdTomato mouse line (*white*) probed by ISH for Dlk1 (*blue, ISH raw signal in left panels*). Segmented Dlk1^high^-expressing cells define a distinct area of the medio-ventral striatum (*blue, segmented high-expressing SPNs in right panels*). **(L)** Within the high Dlk1-expresing area of the medio-ventral striatum Dlk1 (*blue*) is expressed by Oprm1-Cre:tdTomato positive (*white, full arrowhead*) as well as Oprm1-Cre:tdTomato negative SPNs (*empty arrowhead*). **(M-N)** Quantification of Crym positive cells (*magenta*) within the high Crym-expressing area, Gpr155-positive (*green*) cells within the high Gpr155-expressing area and Dlk1-positive (*blue*) cells within the high Dlk1-expressing area. Quantifications are expressed as percentages of Oprm1-Cre:tdTomato positive SPNs **(M)** and percentages of Oprm1:tdTomato negative cells **(N). (O)** Model of the spatial definition of the striatum by the selected spatial markers in the anterior striatum (*upper row*) and posterior striatum (*lower row*). Subregions defined by expression of Crym^high^ (*magenta*), Gpr155^high^ (*green*) and Dlk1^high^ (*blue*) are shown separately (*left*) and as a merged image (*right*). Abbreviations: (AS) anterior striatum, (PS) posterior striatum. Scale bar: B 200 μm, C 1mm, D-E_1_ 200 μm, F 10 μm, G-H_1_ 200 μm, I 10 μm, J-K_1_ 200 μm, L 10 μm.

To directly visualize the distribution of the Col11a1+ cells in striatum, we performed in situ expression mapping together with the patch marker Sema5b in tissue from adult Oprm1:tdTomato mice (Fig. 4C-D). We found that Col11a1 was not expressed in Oprm1:tdTomato+ cells, neither in patch SPNs (i.e. Oprm1+ and Sema5b+) nor in exopatch SPNs (i.e. Oprm1+ and Sema5b-), agreeing with the expression pattern found in the snRNA-seq data. Instead, the Col11a1+ cells (ColSPN) were positive for Sema5b expression, agreeing with the scRNA-seq data, and thereby forming a molecularly distinct SPN category (i.e. Col11a1+, Sema5b+, Oprm1-). Specifically, Col11a1+ cells displayed higher Sema5b expression levels compared to patch Oprm1+ SPNs (Fig. 4D). To conclusively determine whether this candidate Col11a1+ striatal cell type could be neuroanatomically classified as a projection neuron, we performed retrograde labeling of the striatonigral pathway in combination with in situ expression mapping for Col11a1. We found that Col11a1+ cells in striatum were positively labeled following injection of retrobeads into the substantia nigra (SN) (Fig. 4E-G). From this we conclude that the Col11a1+ cells in striatum represent a novel SPN population with striatonigral projections.

### A Spatiomolecular Code in the Striatum

Since the broad spatial organization of the striatum has been primarily based on the topography of corticostriatal projections, we investigated whether it was possible to identify spatially segregated gene expression patterns in our snRNA-seq dataset. We observed that a number of markers displayed either graded or enriched expression across several of the cell-type specific clusters in the t-SNE visualization (Fig. 5A). Interestingly, some markers displayed opposing expression patterns in the t-SNE plot, indicating discrete and potentially biologically relevant patterns. When we investigated the in situ expression of some of the putative spatial markers (e.g. Col6a1, Coch, Cnr1), we found that the in situ signal displayed clear spatial segregation in the dorsal striatum (Fig. 5B). The observed spatial segregation of these markers, evident across the mediolateral axis of the striatum, furthermore corresponded to an axis of marker enrichment also in the t-SNE plot (Suppl. Fig 9). From the snRNA-seq data, we identified a number of candidate markers that can be used to establish a spatiomolecular annotation of the striatum: Wfs1, Col6a1, Coch, Cnr1, Crym, Gpr155, Gpr139, Nefm, Syt10, Rreb, Kctd12, Cpne6, Pde1a (Fig. 5A-B, Suppl. Fig 9).

Similar to the patch-matrix markers, we mapped the in situ expression of available genes in the Marmoset Gene Atlas to determine the conservation of the spatial signals. We found that the candidate spatial markers (Crym, Wfs1, Dlk1, Syt10, Cnr1, Cpne6, Nefm) displayed a conserved spatial expression pattern in the caudate and putamen of the marmoset brain (Fig. 5C).

To further map in detail the tissue expression pattern for some of the candidate spatial markers, we mapped the in situ expression for three candidate markers (Crym, Gpr155, Dlk1) in the dorsal striatum in two different anteroposterior segments in coronal sections from Oprm1:tdTomato mice. We found that the expression of Crym defined a medial region of striatum in anterior as well as posterior sections (Fig. 5D-F). To investigate whether the spatially segregated expression of Crym was a general feature of SPNs or was restricted to only one SPN subtype, we quantified Crym expression in tdTomato+ and tdTomato-striatal cells (Fig. 5M). We found that Crym expression was found in 95.1 % of the Oprm1-Cre:tdTomato SPNs localized in medial striatum, which also was supported by the detection of Crym in all major D1+ as well as D2+ clusters in the snRNA-seq data (18% of D1+ nuclei are Crym+, 19% of D2+ nuclei are Crym+).

We further mapped the in situ expression of Gpr155, and we found that it was restricted to a spatially defined central and lateral domain, which was spatially distinct compared to the Crym+ medial subregion (Fig. 5G-I). Similar to Crym, Gpr155 was found to be expressed in both tdTomato+ and tdTomato-striatal cells in the Gpr155+ striatal region (Fig. 5M-N). Supporting the cell-type independent nature of the spatial code, the expression of Gpr155 was not restricted to D1+ or D2+ nuclei in the snRNA-seq data (73% of D1+ nuclei are Gpr155+, 87% of D2+ nuclei are Gpr155+). The expression of Dlk1 instead labeled a discrete medioventral part of the striatum (Fig. 5J-L), which was again spatially distinct from the Crym+ and Gpr155+ domains. Mapping of Dlk1 in situ expression at the cellular level supported the finding that these markers represent a spatial code (Fig. 5M-N). When we combined the distribution of segmented cells with expression of one of the spatial markers (Crym, Gpr155, Dlk1), we found that these could form a model representing the spatiomolecular code of the striatum (Fig. 5O).

In summary, we propose that the spatial code is independent of the classical cell-type specific code, and it is possible using spatial markers to map the dorsal striatum into spatiomolecular subregions.

## DISCUSSION

We have in this study mapped the identity of the striatal projection neurons (SPNs), focusing on establishing the molecular heterogeneity of patch and matrix SPNs, in order to reveal the logic underlying the striatal organization in terms of a spatiomolecular code. This study provides the first demonstration of molecular codes that divide the dorsal striatum into three main levels of classification: a) a spatial organization, b) a patch-matrix organization, and c) a cell-type specific organization. A key observation is that the markers displaying spatially segregated expression (the spatial code) are not confined to a single SPN subtype, but rather represent a shared code found in all SPNs that are localized within a defined spatial domain. We therefore propose that the spatial code represents a different level of classification, in addition to the cell-type specific gene expression code, and that this spatial code can be applied to define the primate caudate and putamen subregions. We furthermore found molecularly discrete SPN subtypes beyond the classical D1-D2 classification, including Oprm1+/Sema5b+ exopatch SPNs and Col11a1+ striatonigral SPNs, in addition to new cellular markers for defining the patch-matrix division. We anticipate that these SPN subtypes will display unique properties in terms of input-output connectivity and specific function during behavior, although this remains to be studied.

### Molecular Anatomy of the Patch-Matrix Division

Striatal patches have neuroanatomically been defined as part of the limbic circuitry, primarily based on the corticostriatal inputs from frontal cortex in combination with a defining projection to dopaminergic neurons in SNc. In contrast, the matrix compartment has been defined as a part of sensorimotor circuit, with preferential inputs from sensorimotor cortex, and outputs to the SNr. Surprisingly, a recent study could not confirm the previously established model of a specialized input organization from cortical layers onto striatal patches^23^. It is possible that this discrepancy could be a result of the use of rabies tracing, in contrast to more traditional non-genetic tracers, in combination with genetic labeling of both patch as well as exopatch neurons. Here we show that it is possible to differentiate between the patch and matrix compartment using cellular markers, thereby establishing a discrete molecular code that can be employed to develop genetic targeting strategies.

The spatial boundaries between patch and matrix compartments have up to now relied on immunostaining against MOR, resulting in ambiguous definitions at the cellular level. Genetic and cellular definitions of neuron identity allow for a precise and systematic classification of the striatal circuit. Genetic targeting of SPNs based on BAC transgenic mice (Sepw1-NP67 Cre line)^23^ or developmental birth dating of SPNs (Mash1-CreER)^31^ has allowed for the first time the genetic labeling of cells in the patch compartment. Such approaches, including approaches of visualizing striatal patches using Pdyn-GFP^32^ or Nr4a1-GFP^33^ mice are pivotal to understanding the functional specialization of the patch-matrix compartment. Recent studies have imaged the activity of SPN subtypes during motor and choice behavior^5,34^, including the activity of putative patch SPNs^31^, and the complexity of the activity patterns found in vivo represent a challenge in understanding how the dorsal striatum orchestrates action selection and motor behavior.

In order to genetically identify the patch compartment and the striatal cells that form the characteristic dense MOR immunostaining, we generated an Oprm1-Cre mouse line that allowed us to visualize with cellular resolution the MOR-positive striatal cells

We found a number of candidate markers to distinguish the patch versus matrix compartments from our snRNA-seq data. To our knowledge, these patch markers allow us for the first time to define the patch identity at the cellular level. Interestingly, when we investigated whether our candidate patch-markers also defined spatially segregated expression in the dorsal striatum of a non-human primate, we found that all the available markers showed a distinct spatial expression in the marmoset caudate and putamen, which is characteristic of the patch organization.

When we in detail mapped the in situ expression of candidate patch-matrix markers, we found that Sema5b and Id4 expression could define at the cellular level the patch versus matrix identity of SPNs. We found that patch SPNs are labeled with Oprm1+/Sema5b+ expression whereas matrix SPNs (D1+ as well as D2+) are Id4+. In addition to the Sema5b-Id4 division, we found a number of other markers that support this patch-matrix division (patch: Mfge8, Kremen1, Pde1c, Lypd1; matrix: Sgk1, Sv2b, Epha4). In summary, these markers represent a code that captures the patch-matrix identity at the cellular level. It is interesting to note that the expression of these candidate patch markers in the striatal was not identical all markers, suggesting a molecular differentiation of patch SPNs in different striatal subregions. For example, some patch markers (Crhr1, Lypd1, Tshz1) showed a classical patch labeling pattern, including labeling of the MOR-dense subcallosal streak, whereas other patch markers (Nnat, Rab3b) labeled patches in medial or central striatum (Suppl. Fig 6).

The molecular relationship between the classical patch SPNs and the exopatch SPNs has remained unclear. A recent study reported that patch and exopatch SPNs displayed similar genetic, neurochemical and electrophysiological properties^23^. We found that exopatch cells share some of the molecular code with patch SPNs (i.e. Oprm1 expression), but are also clearly distinct from both patch as well as matrix SPNs in terms of expression of for example Sema5b. The combination of the three markers Oprm1/Sema5b/Id4 can be used to classify the patch-matrix division including exopatch SPNs.

A recent scRNA-seq study identified graded gene expression in SPNs and clustered striatal cells into subtypes based on identification of new markers (e.g. Pcdh8, Dner)^36^. In our study we found that Pcdh8 and Dner expression reflected the identity of cell-type specific clusters. Pcdh8 expression was primarily found in the Col11a1+ SPNs and the Pvalb+ interneurons, whereas Dner was enriched in Col11a1+ SPNs and several interneuron types. Furthermore, the gene expression of some of the markers that we defined as spatially segregated have previously been categorized as continuous transcriptional gradients in subtypes of striatal cells (Dner, Cnr1, Crym, Wfs1). These gradients in the expression level were described to form opposing gradients (e.g. Crym-Cnr1), which is similar to the expression pattern of spatially segregated markers in our snRNA-seq data. Based on our analysis, we conclude that these expression patterns do not in themselves define subtypes of SPNs but instead indicate the spatial identity of SPNs.

Two recent scRNA-seq studies mapped the expression profile of single neurons throughout the forebrain, including the striatum, resulting in the identification of the major SPN subtypes studies^30,37^. These studies demonstrated the molecularly diversity of the striatum, revealing new markers for striatal cell types. One study identified primarily gene expression differences in the dorsoventral axis, mapping two D1+ SPNs subtypes and two D2+ SPNs subtypes to either dorsal or ventral striatum, respectively, and also described one patch group (markers: Drd1, Lrpprc, Tshz1) and one matrix group (markers: Tac1, Wnt2, Gng2)^37^. The main markers for the patch-matrix division was based on patch-enriched genes, either previously known enriched markers such as Tac1, as well as new markers (e.g. Tshz1, Asic4, Necab1). The expression of Asic4 was reported to be enriched in D2+ patch SPNs, and Necab1 expression in D1+ patch SPNs^30^. We observed that some of the proposed markers for patch SPNs (Asic4 and Necab1) were not defining markers of the patch clusters in our snRNA-seq data. We found that Asic4 expression predominantly labeled D1+ SPNs in both patch and matrix, whereas Necab1 expression was enriched in the D2+ SPNs, although it was also found in D1+ SPNs and the Col11a1+ SPNs. Technical differences in the RNA-seq methodology including in the sensitivity between snRNA-seq and droplet microfluidics scRNA-seq could potentially explain differences in marker detection. Ultimately, it will be valuable to perform a considerably deeper and more extensive scRNA-seq characterization of SPNs across the entire striatal volume, including extensive comparative analysis with other species, to define in detail the molecular code that can describe all the cell types and subregions of the striatum.

### The Spatiomolecular Code Defines Striatal Subregions

What does the spatial code represent in terms of function? It is possible that the spatiomolecular code overlaps with specific input-output patterns of the striatal subregions. For example, the topography of the corticostriatal projections could map onto discrete molecular domains in the striatal volume. A number of studies have mapped the spatial segregation of the corticostriatal terminals in the striatum^38,39,40,41^, but how this connectivity maps onto discrete gene expression domains remains unknown. The corticostriatal inputs can be used to subdivided the striatum (defined as intermediate CP, CPi) into four core domains that are segregated in the dorsoventral and mediolateral axes^39^. Mapping this connectivity map onto our spatiomolecular annotation, reveals the dorsomedial domain (CPi.dm) as the best corresponding to the domain captured by Crym expression, whereas the ventromedial connectivity domain is best represented by the expression of Dlk1.

An interesting perspective is that the spatial code can also reflect a specialization in signaling pathways, since for example the spatial markers Gpr155 and Gpr139 are both G-protein-coupled receptors (GPCRs), although the identity of their ligands remains unknown. In that context, other signaling molecules or adhesion complexes could form spatial domains that support the functional and/or structural organization of the basal ganglia. Since the developmental trajectory of striatal progenitors can give rise to the different SPN types^42^, it is possible that the combination of a developmental history and accompanying establishment of the connectivity pattern ultimately are reflected in a molecular profile that can define spatial boundaries in the adult striatum.

Based on our mapping of striatal snRNA-seq and in situ expression data, we propose that the cellular classification (SPN subtypes) is nested within higher-order classification of space (tissue subregions). Importantly, this classification scheme is possibly conserved in primates based on the available data in the marmoset brain, pointing to an evolutionary conserved organization of the striatal cell types and tissue architecture.

In addition to the cell-type specific classification, the classification of striatal tissue subregions has been important in guiding the definition of functionally segregated regions in terms of their role in shaping behavior. The spatiomolecular map can ultimately help to integrate molecular definitions of cell types and subregions (tissue space) with an input-output matrix, and to describe this in relation to specialized functions. The spatiomolecular code can therefore be the key to perform a systematic analysis of the striatal circuitry – by establishing experimentally reproducible targeting of discrete SPN subtypes found in spatially defined domains. Even if gene expression is not strictly binary, it is still possible to use information on gene expression to experimentally define striatal subregions and neuron subtypes. Use of intersectional approaches to define cell type and spatial annotation within the striatum can help develop a systematic and reproducible experimental dissection of the role of cell types and subregions of the striatum.

A number of interesting questions remain regarding the spatial codes: do these represent discrete connectivity profiles, or some specific signaling pathways, and are these established during development? And are there mechanisms to maintain those signals in the adult? Ultimately, it will be important to understand the functional properties of the proteins that generate the spatial code. Neuroanatomical models have been central to developing theories on the function of neuronal circuits, and it is therefore important to establish systematic maps that integrate information on the position, identity and connectivity of neurons. Increasing knowledge on the diversity of neuron types and how molecular information can be used to map cells and tissue in a common reference framework, will guide efforts to map function onto the circuit structure.

## ACKNOWLEDGMENTS

We are grateful to Jonas Frisén for providing access to the FACS facility and Sarantis Giatrellis for operating the FACS; Rickard Sandberg for advice on RNA-seq; Maggie Yeung for advice on the nuclear isolation protocol; Maria Kasper for advice on in situ hybridization; the National Genomics Infrastructure at Science for Life Laboratory for sequencing services; the Eukaryotic Single-Cell Genomics core facility at Science for Life Laboratory for use of equipment. We acknowledge the Allen Brain Atlas and the Marmoset Gene Atlas for generously providing the publicly available in situ hybridization data.

This work was supported by the Swedish Brain Foundation (Hjärnfonden), the Swedish Research Council (Vetenskapsrådet, project 2012-02049), KI doctoral funding (KID) for A.M. and O.T. K.M. was supported by a donation from the William K. Bowes, Jr. Foundation to KI.

## METHODS

### Animals

All procedures and experiments on animals were performed according to the guidelines of the Stockholm Municipal Committee (approval number N166/15). Adult male and female 2-4 months old mice were used: tdTomato Cre reporter (Ai14 line; Jax Stock No: 007914), Vgat-Cre (Jax Stock No: 028862), H2B-GFP Cre reporter^43^, and wild type mice (C57BL/6; Charles River). Oprm1-2A-Cre mice were generated by inserting a T2A-Cre construct into the fifth exon of the Oprm1 gene (Ensembl ENSMUSG00000000766) and replacing a TAA stop codon by homologous recombination in C57BL/6 ES cells (Cyagen Biosciences Inc).

### Tissue dissection and RNA sequencing

For single nuclei sequencing (snRNA-seq), we used striatal tissue from two transgenic mouse lines. Transgenic mice expressing Cre-recombinase in Vgat-positive cells (Vgat-Cre mice) and Oprm1-positive cells (Oprm1-Cre mice) were crossed with mice homozygous for Cre-dependent histone-B associated GFP (H2B-GFP) to obtain mice that express GFP in the nuclei of Vgat-positive neurons (Vgat:H2B-GFP mice) or Oprm1-positive neurons (Oprm1:H2B-GFP mice). The animals were killed with an overdose of isoflurane and the fresh brains were rapidly extracted and immersed in ice-cold ACSF solution. The brains were immediately sliced into 300μm thick tissue sections (around the Bregma +0.62 coordinate) in ice-cold ACSF using a vibratome (VT1200S, Leica). Tissue slices were then submerged in ice-cold Leibovitz’s L-15 medium (Gibco) in a petri dish and the striatum was dissected out and stored in 1 ml ice-cold Leibovitz’s L-15 medium containing 1μl SUPERase RNase inhibitor (20 U/μl, Thermo Fisher Scientific). The neuron nuclei were isolated using a nuclear isolation protocol^25^. The tissue was homogenized in 2 ml lysis buffer using a Dounce tissue grinder (7ml, VWR). This suspension was supplied with a 1.8M sucrose solution (4 mL), homogenized and then added onto a sucrose cushion (2ml) in a 10ml Ultra-Clear centrifuge tube (Beckman Coulter). The nuclei were then separated from the tissue by centrifugation (26.500xg at 4°C for 1.5 hours). The supernatant was discarded and the nuclei pellet was resuspended in 500μl Nuclear resuspension buffer. Single nuclei were isolated using Fluorescence-Activated Cell Sorting (FACS) by their GFP emission profile, and sorted into 384 well-plates containing 2.3μl ice-cold lysis buffer. Plates containing nuclei were immediately frozen on dry ice and stored on −80°C until further processing. cDNA libraries were produced and sequenced using a Smart-seq2 protocol^26^. Sequencing of the single-nuclei libraries was performed using Illumina HiSeq 2000. The reads were mapped and aligned to the mouse genome (mm10) and gene expression values were calculated as count values for each transcript. Analysis was performed on count values of the exome per nucleus.

### Gene expression clustering and marker identification

The raw snRNA-seq data counts were subjected to a strict quality control using the scater R package (version 1.2.0, https://bioconductor.org/packages/release/bioc/html/scater.html). We analyzed nuclei that expressed a minimum of 2000 genes, and showed a spike expression ratio not deviating more than 3 standard deviations (SDs) and a library size of not lower than 3 SD. Genes were excluded from analysis if they were not expressed in at least 3 nuclei. The remaining nuclei and genes were analyzed using the Seurat R package (versions 1.4.0 and 2.3.3, https://github.com/satijalab/seurat; http://satijalab.org/seurat/). The raw count data were log2 normalized, variance genes were identified by calculating their z-score of log(variance/mean). A principal component analysis (PCA) was performed followed by random sampling with 1000 replicates to determine the significant Principal Components (PCs) in the dataset. A projected PCA was done to increase the gene list and to prevent missing potential marker genes. The resulting list of genes was analyzed for principal components and randomly sampled (1000 replicates). A nonlinear dimensionality reduction (t-SNE, 2000 iterations) was performed on the identified significant PCs (max. 100 genes per PC were allowed in the clustering to prevent one PC from driving the entire clustering). Density-based clustering was performed and markers per cluster were identified based on their differential expression.

### In situ hybridization

In situ hybridization (ISH) to detect expression of genes of interest was performed using RNAscope technology following the manufacturer’s protocol (Advanced Cell Diagnostics, ACD, Hayward, CA). Highly expressed genes (*Tac1, Adora2a, Id4, Sema5b, Crym*) were visualized using the RNAscope Fluorescent Multiplex Assay (cat. no. 320850). For genes with lower expression values (*Oprm1, Gpr155, Gpr139, Col11a1*) the HRP-based RNAscope Fluorescent Multiplex Assay V2 was used (cat. no. 323110). For patch-matrix and cell type specific genes ISH was used on tissue sections from the Oprm1-Cre-tdTomato mouse line. For genes related to spatial distribution ISH was performed on tissue sections from wild type mouse brains (C57BL/6, Charles River). Adult mice were transcardially perfused with 1x PBS followed by 4% paraformaldehyde and post-fixation in 4%PFA overnight. Brains were subsequently cryoprotected in a 15% sucrose solution followed by a 30% solution (in PBS) overnight at 4°C. After cryoprotection, brains were frozen on dry ice and stored at −80°C. Brains were sectioned into 14 μm thin sections on a cryostat (CryoStar NX70 cryostat, Thermo Fisher Scientific) and stored at −80°C until further processing. The tissue sections were subjected to permeabilization steps before incubation with the target probes. The signal was amplified using either the HRP-based technology (visualized by TSA fluorophores by Perkin Elmer) or the RNAscope Fluorescent Multiplex amplification method.

### MOR immunohistochemistry

Immunohistochemistry for the mu opioid receptor (MOR) was performed in Oprm1-Cre:tdTomato mice (n = 3). Brain sections were cut on a vibratome at 50 μm thickness (Leica VT1000, Leica Microsystems GmbH) and were placed in sodium citrate (10 mM Sodium Citrate, 0.05% Tween 20, pH 6) for 2 min for antigen retrieval followed by wash in tris buffered saline solution (TBST) including 0.3% Triton. After antigen retrieval, blocking was performed with 5% donkey serum in TBST for one hour at room temperature (RT). Sections were then incubated with the MOR primary antibody (1:1000 in TBST, Neuromics, cat. no.RA10104) on a shaker at RT overnight. The second day sections were washed with TBST (10min, RT) followed by a 2-hour incubation with the HRP conjugated reagent (1:1 in TBST, HRP Polymer Conjugate Rabbit Primary, Life Technologies, LOT:1709065A) at RT on a shaker. Brain sections were washed with TBST (10min, RT) before incubated with the Fluorophore Tyramide Amplification Reagent (1:200 in amplification diluent, TSA Plus multi-fluorophore detection kit, Fluorophore: Fluorescein, PerkinElmer, NEL753001KT). Sections were washed overnight in 1x PBS at RT. The sections were washed in TBST (10 min, RT), and then with 1x PBS (10 min, RT). Sections were mounted on cover glass with SlowFade Gold antifade reagent (Life Technologies, S36936).

### Tissue clearing

To visualize Oprm1-Cre:tdTomato+ neurons in the whole striatum 3 months old Oprm1-Cre-tdTomato mice were transcardially perfused with 30 mL 0.1 M phosphate buffered saline (PBS) solution (pH 7.6), followed by 100 mL of 4% PFA in 0.1 M PBS (pH 7.6). The brain was extracted from the skull and post-fixed overnight in 20 mL of 4% PFA at 4°C. After rinsing in 0.1 M PB (pH 7.6), coronal sections of the left and horizontal sections of the right hemisphere (750 um) were prepared using a vibratome (Leica VT1200S). Slices containing the striatum (4 coronal, 4 horizontal) were selected and further processed for CUBIC tissue clearing^24^. In summary, slices were incubated in CUBIC1 solution (25% urea, 25% *N,N,N′,N′*-tetrakis-(2-hydroxypropyl)ethylenediamine and 15% Triton X-100) at room temperature for 4 days under continuous agitation. After 3 washes in 0.1 M PB (pH 7.6) samples were immersed in CUBIC2 solution (50% sucrose, 25% urea, 10% 2,2′,2″-nitrilotriethanol, and 0.1% Triton X-100) and left shaking overnight before imaging. Slices were mounted on Superfrost glass (Thermo Scientific) using CUBIC2 solution and covered with 1.5 mm cover glasses.

### Image acquisition and analysis

All confocal images were taken using a Zeiss 880 confocal microscope. For CUBIC cleared Oprm1-Cre-TdTomato sections were imaged using a Plan-Apochromat 10x/0.45 M27 objective (imaging settings: frame size 512x512, pinhole 1AU, bit depth 8 bit, speed 7, averaging 2). For ISH overviews a Plan-Apochromat 20x/0.8 M27 objective was used (imaging settings: frame size 1024x1024, pinhole 1AU, Bit depth 16 bit, speed 6, averaging 2). For ISH close up imaging a Plan-Apochromat 63x/1.40 Oil DIC M27 objective was used (imaging settings: frame size 1024x1024, pinhole 1AU, Bit depth 16 bit, speed 6, averaging 4). Confocal stacks were stitched using Zen Blue software (Zeiss) before importing the into Imaris software (Bitplane, version 9.2). Images from the Oprm1-Cre-tdTomato mouse brains immunostained with MOR, were acquired on a Leica DM6000B fluorescent microscope with a 16-bit depth resolution Hamamatsu Orca-FLASH 4.0 C11440 digital camera, at 10X (four striatal coordinates: 1.1, 0.9, 0.4, 0.1 mm from bregma) and were stitched using the WholeBrain software^44^. The tdTomato labeled neurons were segmented and registered in a standardized mouse brain reference atlas, using a custom written code in R statistics provided by http://wholebrainsoftware.org/. A dataset per section was generated, in which each row represented one segmented neuron and columns carried information about the individual animal number, position from bregma (z position), X and Y position in the brain section, position in the right or left hemisphere. The ontology for each segmented cell was defined using the Allen mouse brain reference atlas annotations. For annotation to patch or matrix compartments, regions of interest (ROIs) were manually drawn around MOR-dense patches (ImageJ). The spatial information from the ROIs was added to the WholeBrain datasets using R. The surface of patches (mm^2^) was calculated by using the ROI Manager in ImageJ (Fiji^45^), whereas the surface of their respective whole striatal areas (mm^2^) was calculated using the openbrainmap interface (http://openbrainmap.org)^44^. For CUBIC cleared sections, tdTomato expressing cells were automatically detected using the *spots* function. Signal detection radius was set at 13 μm. Striatal tdTomato expressing neurons were selected after manually drawing the striatum region 3D borders using the function *surface*. At this point the 3D coordinates of each tdTomato expressing cell was exported for further analysis in Matlab. For ISH quantification tdTomato expressing neurons were automatically detected using the *spots* function, while tdTomato negative neurons were detected manually using the *spots* function on the DAPI channel. In both cases the detection radius was set to 13 μm. In order to have a standardized method to determine ISH positive neurons, RNA expression was quantified by mean intensity within each spot spheric volume on the channel related to the ISH probe. At least 5 negative controls were manually chosen for each probe. At least 100 tdTomato-expressing cells were analyzed for each probe. The threshold for expression was set to the mean plus twice the standard deviation of the negative controls. The validity of this method was tested manually for each probe. Images showing in situ hybridization from the Allen Brain Institute (http://mouse.brain-map.org/) and the Marmoset Gene Atlas (https://gene-atlas.brainminds.riken.jp/) were adjusted for contrast and brightness to enhance the in situ signal using Fiji.

### Striatal patch-exopatch spatial classification

3D co-ordinates of the tdTomato expressing neurons were analyzed with custom written scripts in Matlab. A covering surface was calculated using the detected coordinates, and identical number of simulated cells were generated with random coordinates in the same 3D space. The 5 closest neighbors of each tdTomato neuron and randomly simulated neuron was identified in a spherical space with a radius of 200 um which in every case resulted in more than 5 neighbors. The distance to these neighbors was measured and averaged. Histogram of the average distance values of tdTomato neurons and that of the simulated cells was overlay-compared. The threshold for patch-exopatch spatial classification was defined by the 95 percentile of the Gaussian-distribution of the random-generated neurons (likelihood of misclassifying a cell as an exopatch neuron was kept below 5%). Applying the individual threshold for each striatal brain slice, the original tdTomato positive neuronal coordinates were divided into patch and exopatch groups. Patch-exopatch automated spatial classification was performed in the undivided coronal striatal slices. The sections were first divided into four blocks, each representing 25% of the medio-lateral slice width. Separate covering surfaces were calculated onto each block in order to compute block volume for cell density normalization. Patch and exopatch neurons were separately quantified in each block and neuronal densities were expressed as cells/mm^3^.

### Stereotaxic injections

For injections of red retrobeads (Lumafluor) in the substantia nigra, 3 months old wild type mice were anesthetized with 5% isoflurane and mounted in a stereotaxic apparatus (Harvard Apparatus). 100 nl of retrobeads were injected unilaterally using a glass micropipette at 50 nl/minute with a Quintessential Stereotaxic Injector (Stoelting). Substantia nigra was targeted using the following coordinates: 3.3 mm caudal and 1.5 mm lateral to bregma, and 4.15 mm deep from the dura (2 mice). The glass pipette was left in place for 15 minutes after the injection and before removal from the brain. Mice were killed 4 days post-injection for tissue histology.

## SUPPLEMENTARY FIGURE LEGENDS

**Supplementary Figure 1 (related to Figure 1).**
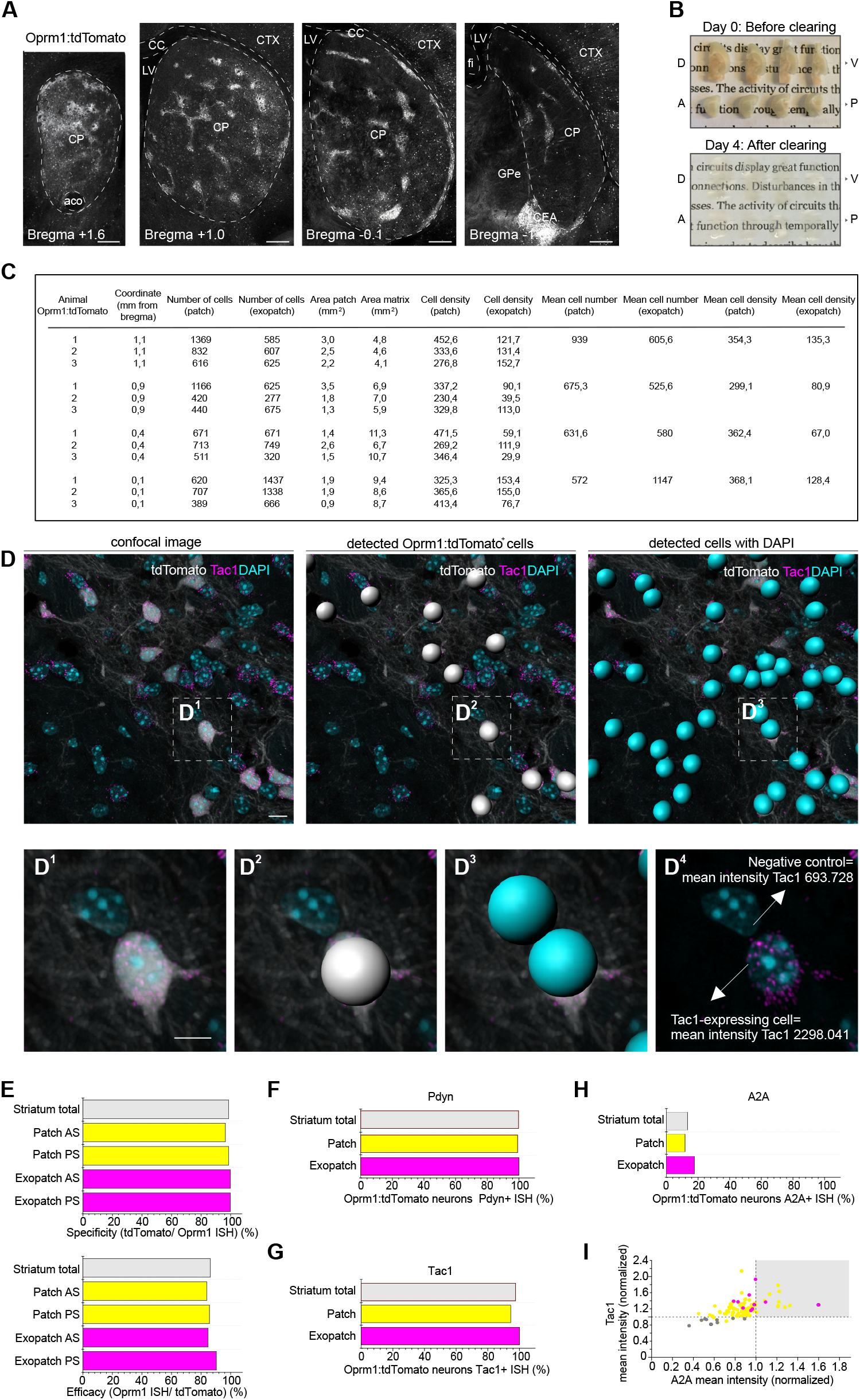
Mapping of Striatal SPNs in Oprm1-Cre:tdTomato mice. **(A)** Representative overviews illustrate the distribution of Oprm1-Cre:tdTomato positive SPNs. **(B)** Four consecutive dorso(D)-ventral(V) 750 μm thick horizontal sections and four anterio(A)-posterior(P) 750 μm thick coronal sections containing the whole striatum are displayed before (*upper panel*) and after CUBIC clearing protocol (*lower panel*). **(C)** Table corresponding to Figure 1B. **(D)** Representative example of ISH probes automated mean intensity detection method. **(D_1_)** High resolution images after ISH (*Tac1 in magenta*) and DAPI staining (*cyan*) on Oprm1-Cre:tdTomato (*white*) slices in Imaris software. **(D_2_)** tdTomato expressing neurons are automatically detected using the spots function (*gray spots*). **(D_3_)** Using the DAPI channel all the DAPI positive cells are detected (*blue spots*). **(D_4_)** Representative example of mean intensity values of the Tac1 probe channel (ISH, *magenta*) within tdTomato negative detected spot (negative control, mean intensity=693.728) and within tdTomato positive SPNs (mean intensity= 2298.041). **(D)** Specificity and efficacy of Oprm1-Cre:tdTomato mouse quantified in the anterior and posterior striatum (AS-PS) within Patch SPNs (*yellow*), within Exopatch SPNs (*magenta*) and in the total Oprm1-Cre:tdTomato positive population. **(E-G)** Quantification of Oprm1-Cre:tdTomato positive SPNs expressing Pdyn **(E)**, Tac1 **(F)** and A2A **(G)** within Patch SPNs (*yellow*), Exopatch SPNs (*magenta*) or total Oprm1-Cre:tdTomato positive population. (*grey*). **(H)** Tac1 (ISH) versus A2A (ISH) expression levels in negative controls (*gray*), in Patch (*yellow*) and in Exopatch SPNs (*magenta*) is visualized as single cell normalized mean intensity. Threshold for expression of each probe was set to the mean plus twice the standard deviation of the negative controls (*dashed line*). Abbreviations: (aco) anterior commissure, (CC) corpus callosum, (CEA) central amygdalar nucleus, (CP) caudoputamen, (CTX) cortex, (fi) fimbria, (GPe) globus pallidus external segment, (LV) lateral ventricle. Scale bar: A: 200 μm, D-D_4_: 10 μm.

**Supplementary Figure 2 (related to Figure 1).**
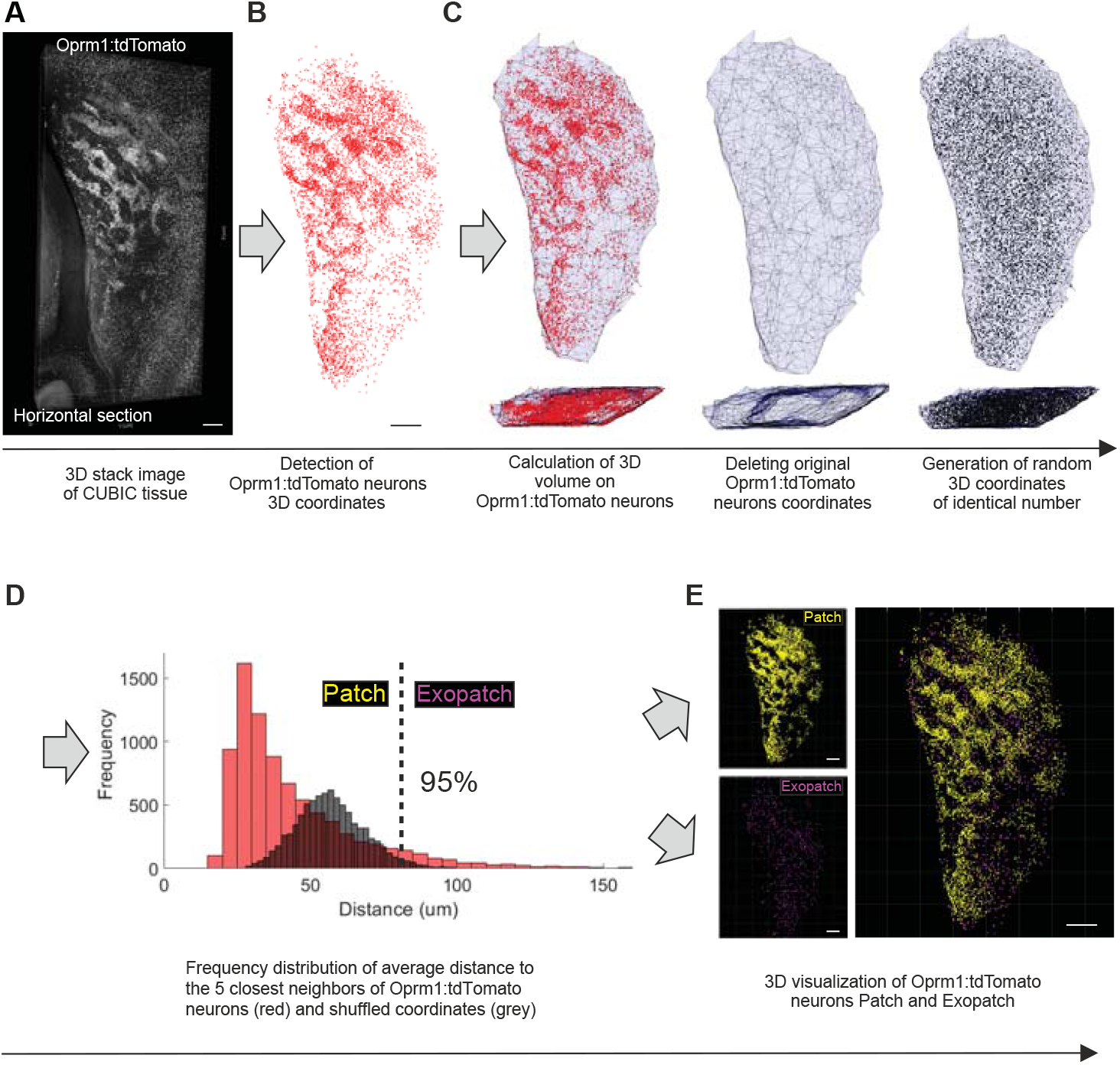
Classification of Patch versus Exopatch SPNs based on automated spatial mapping. **(A)** Representative image of a 750 μm thick CUBIC cleared horizontal section of the adult striatum showing Oprm1-Cre:tdTomato labeled cells. **(B)** 3D coordinates of the Oprm1-Cre:tdTomato positive SPNs (*red spots*) are automatically detected. **(C)** A 3D covering surface was calculated using the detected coordinates (*gray background*) and an identical number of simulated cells were generated with random coordinates in the same 3D space (*black spots*). Top view (upper panels) and side view (bottom panels) of the 3D space analyzed. **(D)** Overlaid histograms of the average distance values of the five closest neighbors to each tdTomato positive neuron (*red*) and randomly simulated neuron (*gray*). The threshold for Patch *versus* Exopatch spatial classification was defined by the 95th percentile of the distribution of the random-generated neurons (*dashed line*). **(E)** Original Oprm1-Cre:tdTomato positive SPNs are classified into Patch (*yellow*, upper panel) and Exopatch (*magenta*, lower panel). Scale bar: A-B, E 200 μm.

**Supplementary Figure 3 (related to Figure 1).**
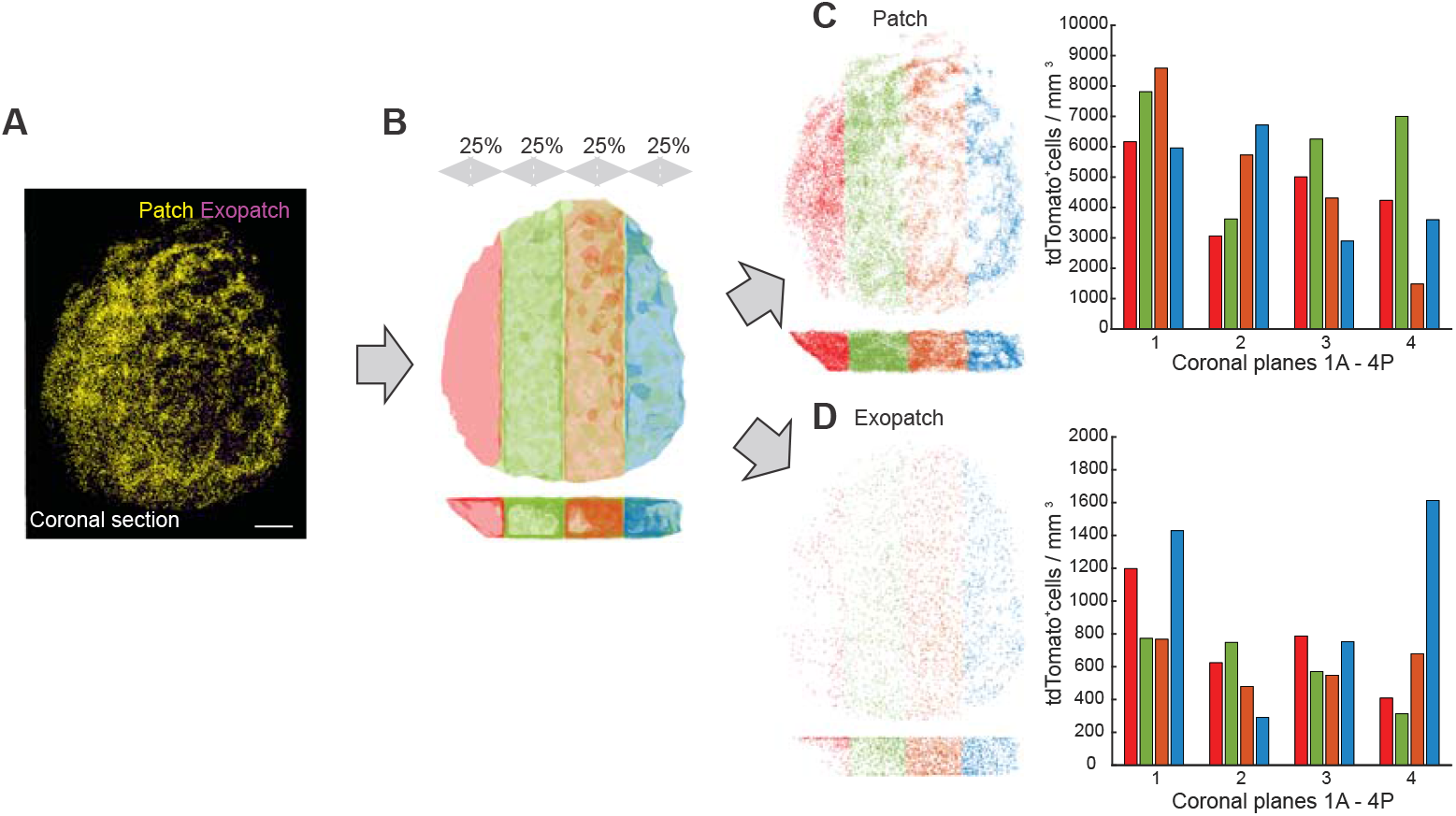
Patch and Exopatch SPNs distribution quantified in the medio-lateral axes of the striatum. **(A)** Representative image after automated spatial classification of Patch (*yellow*) and Exopatch SPNs (*magenta*) in a 750 μm thick coronal striatal section. **(B)** Illustration of a thick coronal slice volume virtually divided into four blocks, each representing 25% of the medio-lateral slice width. **(C-D)** Patch SPNs **(C)** and Exopatch SPNs **(D)** are separately quantified in each block and neuronal densities are expressed as cells/mm^3^.

**Supplementary Figure 4 (related to Figure 2).**
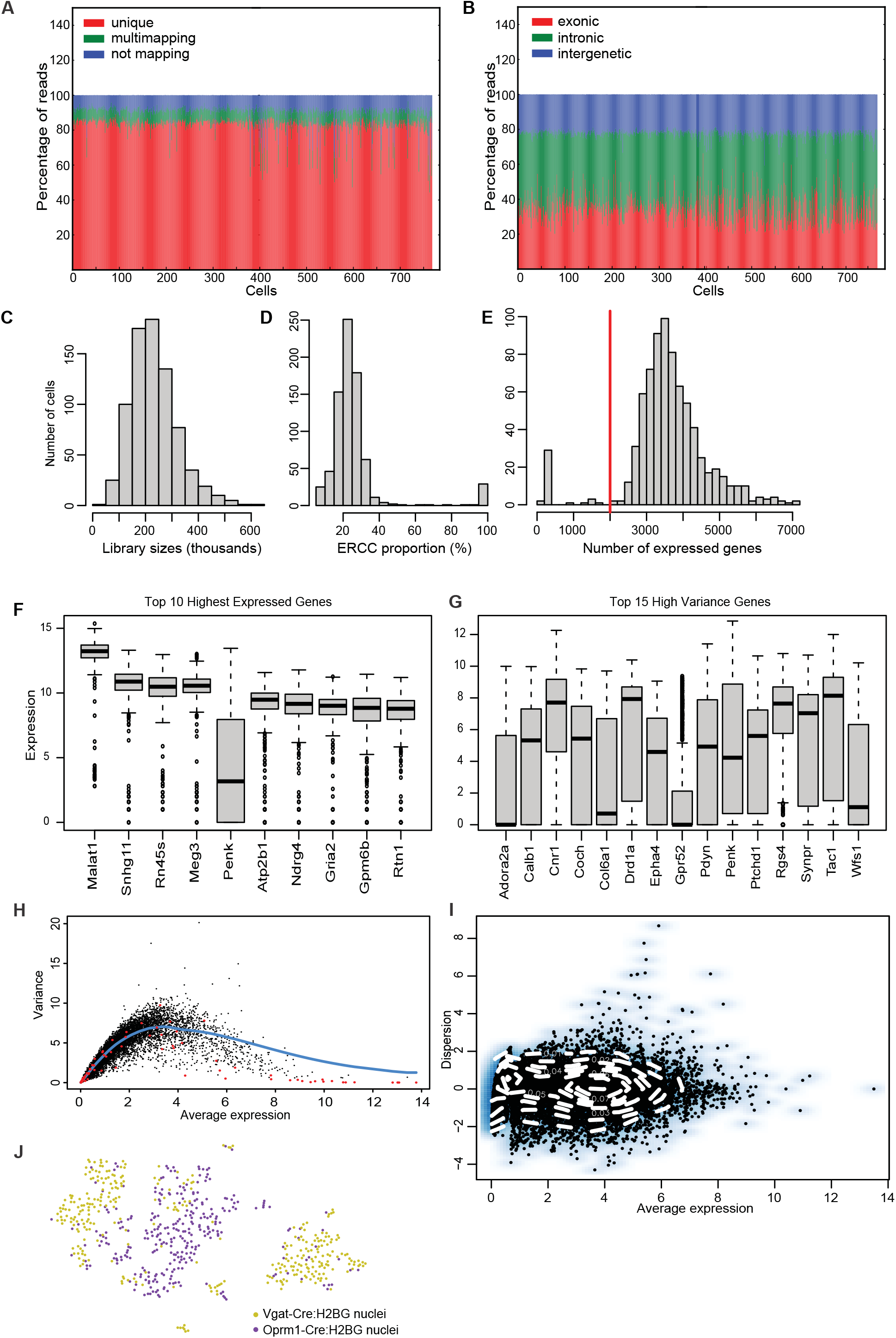
Quality Control Measures of snRNAseq data. **(A)** Percentage of reads aligning to reference genome (*mm10*) over the whole nuclei population (*n=768*). On average, 80% of reads uniquely map to the reference genome (red), while 10% map either to multiple regions (depicted in green), or no region (depicted in blue). **(B)** Fraction of reads aligning within exons (red, average 30%), introns (green, average 50%) and intergenic regions (blue, average 20%). **(C)** Distribution of library sizes over the whole population. **(D)** Ratio of Spike-ins (ERCCs) relative to gene expression. Included in the analysis are nuclei with a ratio below 3 standard deviations above the mean. **(E)** Number of genes expressed in the nuclei. Included in the analysis are nuclei expressing a minimum of 2000 genes (red vertical line). **(F)** Top Ten Highest Expressed Genes for the entire dataset. Gene expression (*y-axis*) is given in normalized log-expression. **(G)** Top 15 High-Variance Genes. Gene expression (*y-axis*) is given in normalized log-expression. **(H)** Variance of each gene plotted against its average count (given as log-expression). The red dots highlight the mean-dependent trend in the technical variance of the ERCC spikeins. The fitted trend of variance is represented by the blue line. **(I)** Identification of highly variable genes. The dispersion of the genes (*variance/log-average expression*) is plotted against the average expression (*normalized log-expression, on x-axis*). Our minimum threshold to be considered highly variable was set at a dispersion of 2 on the y-axis. **(J)** Mouse origin for the isolated nuclei in all clusters in the t-SNE plot (purple dots represent nuclei from the Oprm1-Cre:H2BG mouse line, yellow dots from the Vgat-Cre:H2BG mouse line).

**Supplementary Figure 5 (related to Figure 2).**
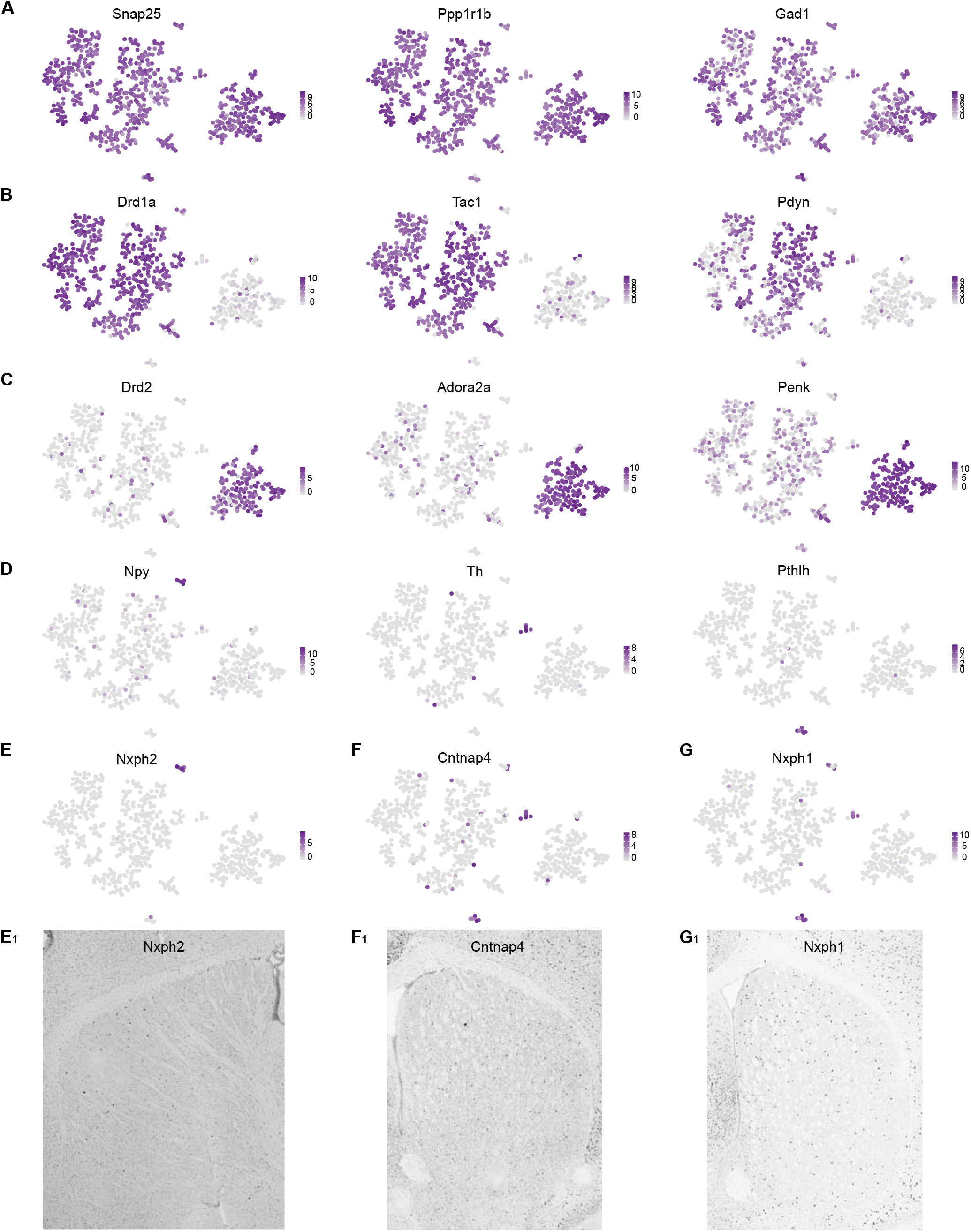
Distribution of cell-type specific marker expression. **(A)** Expression of molecular markers in the t-SNE plot. **(B)** Expression of D1 SPN markers in the t-SNE plot. **(C)** Expression of D2 SPN markers in the t-SNE plot. **(D)** Expression of interneuron markers in the t-SNE plot. **(E)** Expression of Nxph2, a new molecular marker for the Npy interneuron cluster in the t-SNE plot and **(E1)** the corresponding in situ hybridization image **(F)** Expression of Cntnap4, a new molecular marker for the Th interneuron cluster, shown in the t-SNE plot and **(F1)** the corresponding in situ hybridization image. **(G)** Expression of Nxph1, a new molecular marker for Pthlh interneuron cluster, shown in the t-SNE plot and **(G1)** the corresponding in situ hybridization image (all ISH data derived from Allen Brain Atlas mouse brain, Allen Brain Institute).

**Supplementary Figure 6 (related to Figure 3).**
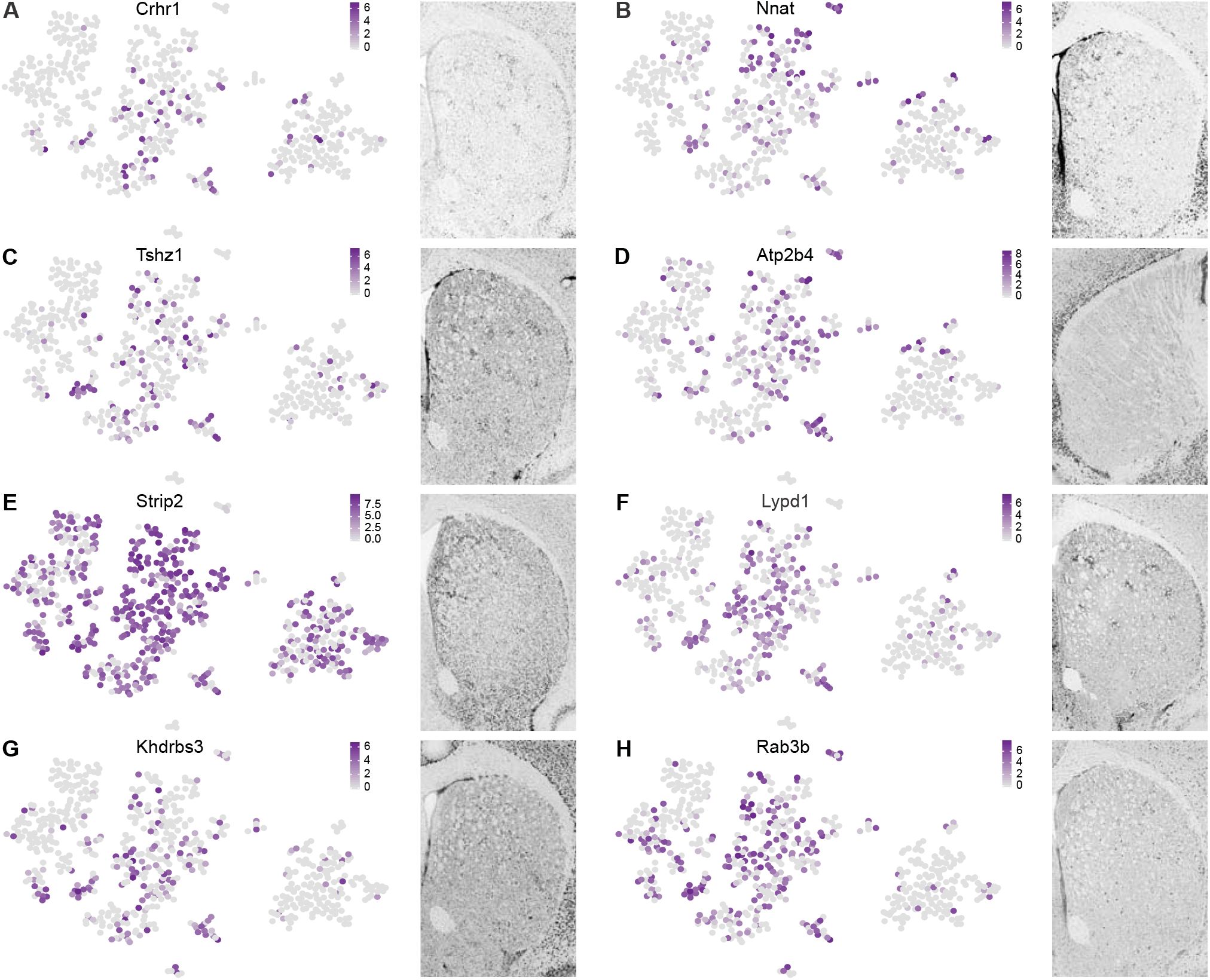
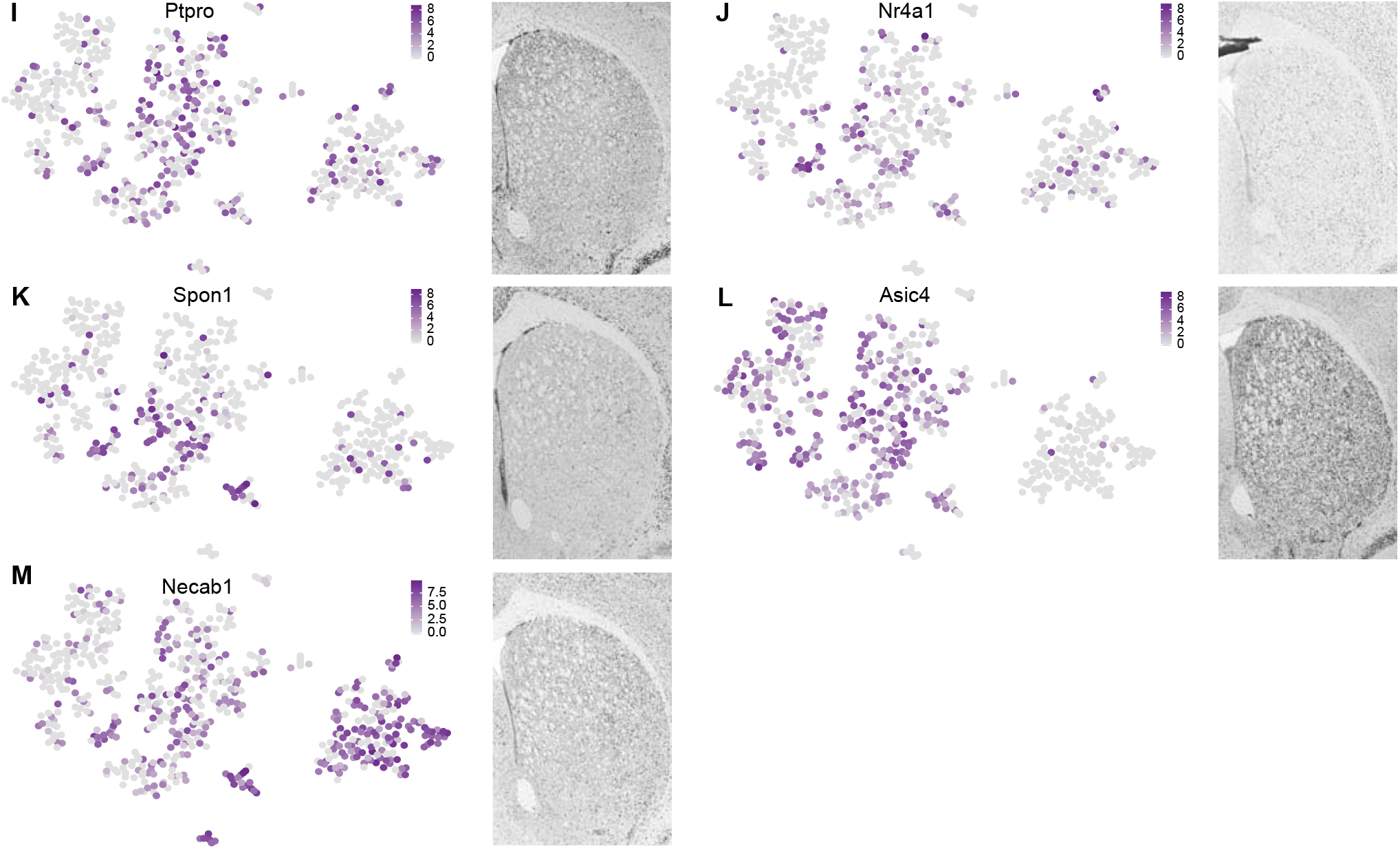
Distribution of striatal patch marker expression. **(A)** Crhr1 expression and corresponding ISH image. **(B)** Nnat expression and corresponding ISH image. **(C)** Tshz1 expression and corresponding ISH image. **(D)** Atp2b4 expression and corresponding ISH image. **(E)** Strip2 expression and corresponding ISH image. **(F)** Lypd1 expression and corresponding ISH image. **(G)** Khdrbs3 expression and corresponding ISH image. **(H)** Rab3b expression and corresponding ISH image. **(I)** Ptpro expression and corresponding ISH image. **(J)** Nr4a1 expression and corresponding ISH image. **(K)** Spon1 expression and corresponding ISH image. **(L)** Asic4 expression and corresponding ISH image. **(M)** Necab1 expression and corresponding ISH image (all depicted ISH data derived from Allen Brain Atlas mouse brain, Allen Brain Institute).

**Supplementary Figure 7 (related to Figure 3).**
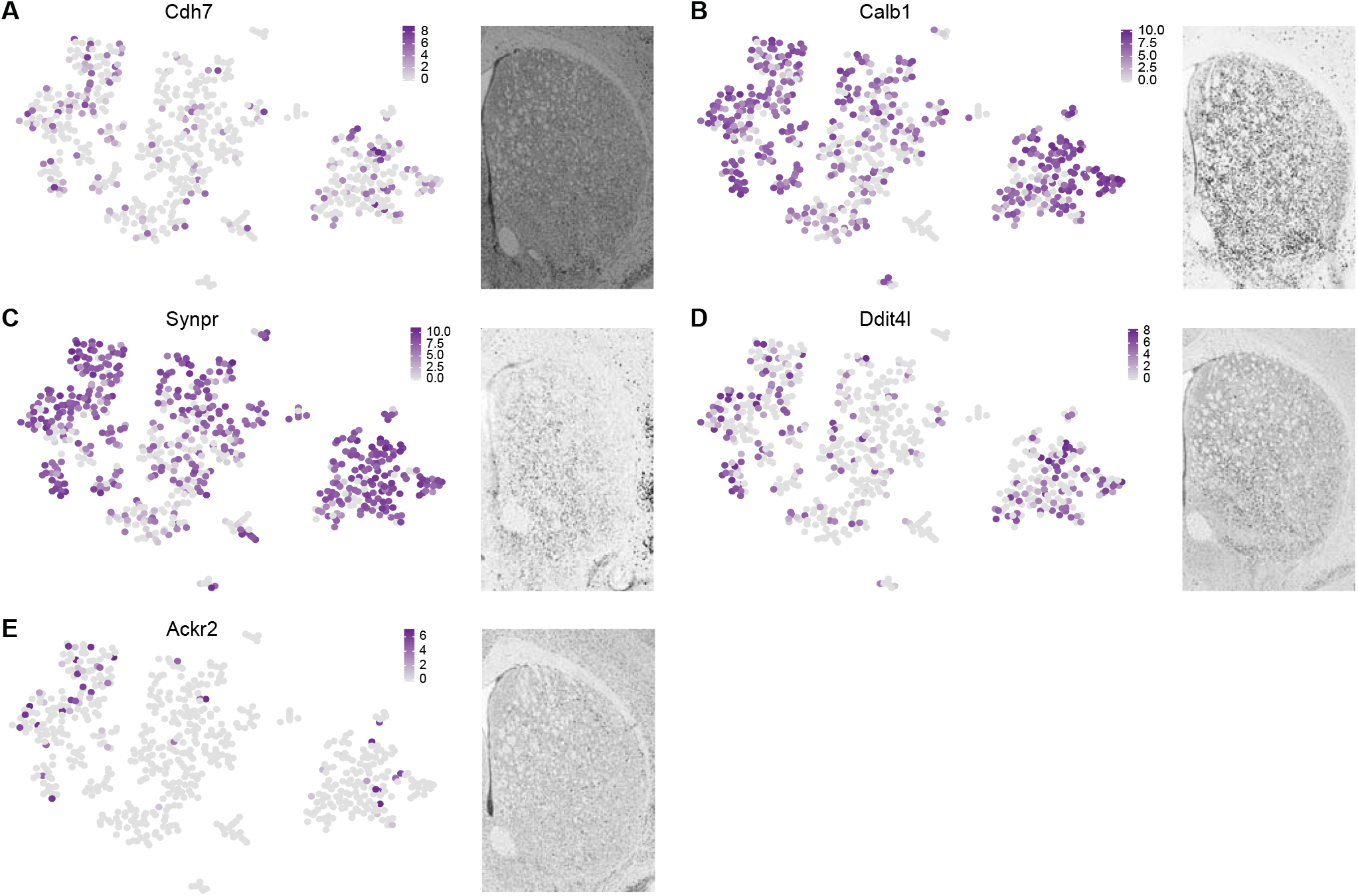
Distribution of striatal matrix marker expression. **(A)** Cdh7 expression and corresponding ISH image. **(B)** Calb1 expression and corresponding ISH image. **(C)** Synpr expression and corresponding ISH image. **(D)** Ddit4l expression and corresponding ISH image. **(E)** Ackr2 expression and corresponding ISH image. (all depicted ISH data derived from Allen Brain Atlas mouse brain, Allen Brain Institute).

**Supplementary Figure 8 (related to Figure 4).**
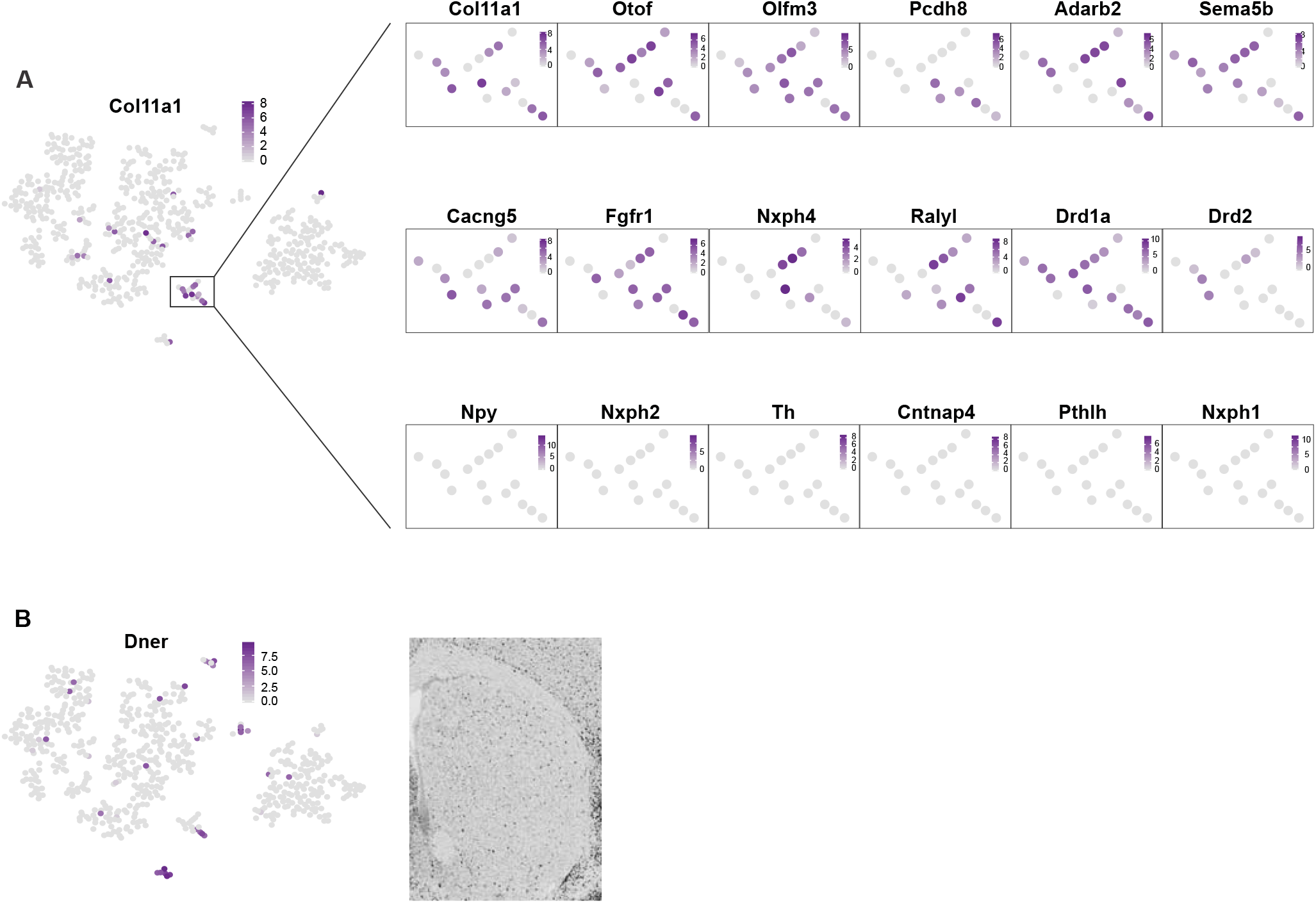
Cell-type specific marker expression in cluster 14 (Col11a1 SPN type). **(A)** Col11a1 expression on t-SNE plot. Cluster 14 in rectangle box. Expression of new Col11a1 SPN type marker genes in cluster 14. Bottom row shows expression of interneuron marker genes in Col11a1 SPN type cluster 14. **(B)** Expression of Dner in the t-SNE plot and corresponding ISH image. (depicted ISH data derived from Allen Brain Atlas mouse brain, Allen Brain Institute).

**Supplementary Figure 9 (related to Figure 5).**
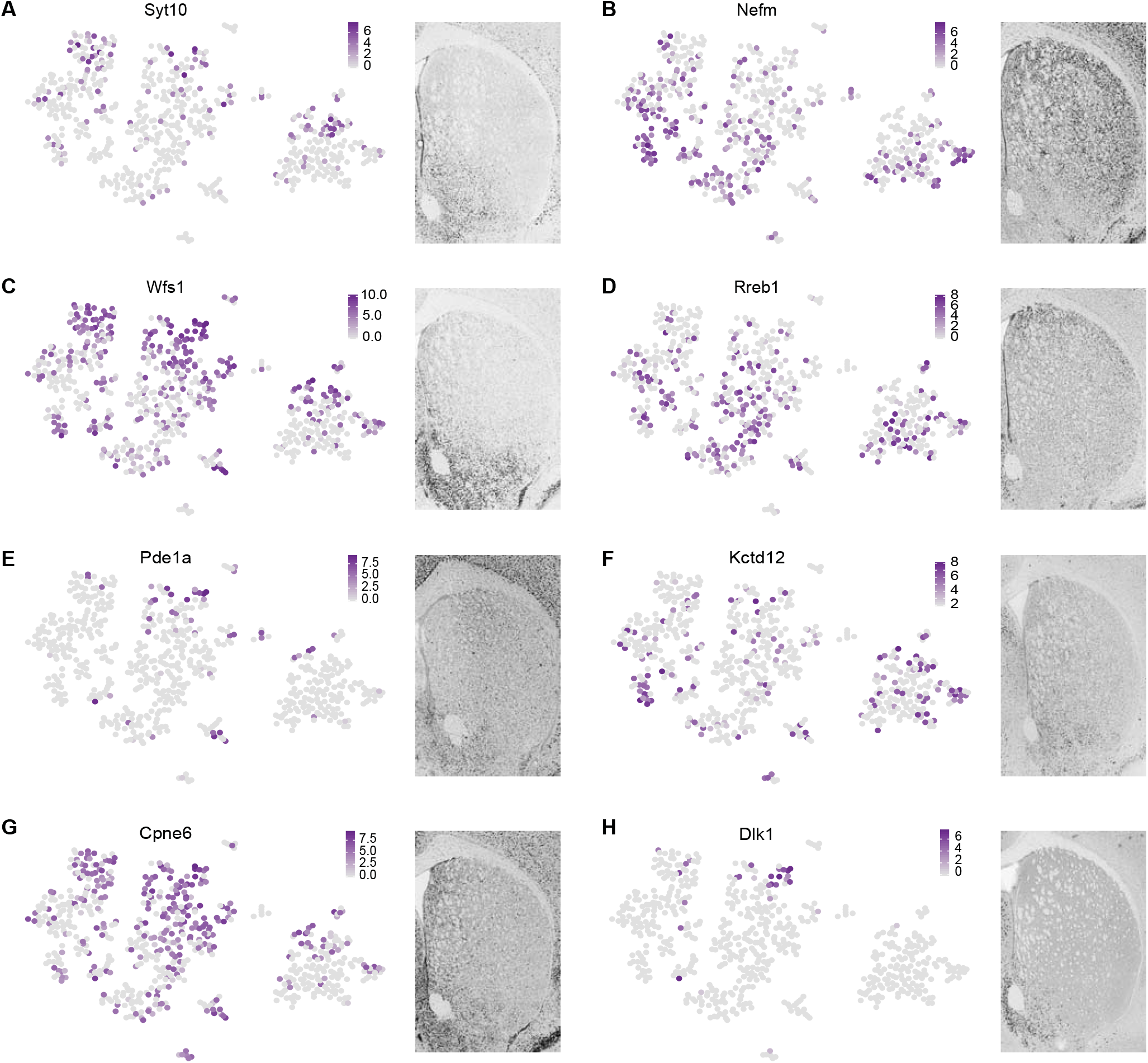
Distribution of spatial marker expression in all clusters. **(A)** Syt10 expression and corresponding ISH image. **(B)** Nefm expression and corresponding ISH image. **(C)** Wfs1 expression and corresponding ISH image. **(D)** Rreb1 expression and corresponding ISH image. **(E)** Pde1a expression and corresponding ISH image. **(F)** Kctd12 expression and corresponding ISH image. **(G)** Cpne6 expression and corresponding ISH image. **(H)** Dlk1 expression and corresponding ISH image. (all depicted ISH data derived from Allen Brain Atlas mouse brain, Allen Brain Institute).

**Supplementary Video 1 (related to Figure 1). Oprm1-Cre:tdTomato positive Patch SPNs and Exopatch SPNs visualized in 3D striatal volume.**

Horizontal 750 μm thick striatal section of Oprm1-Cre:tdTomato mouse after CUBIC clearing is visualized as a 3D volume as well as single optical sections. After automated spatial classification (see methods and Supplementary Figure 2), Patch SPNs (*yellow*) and Exopatch SPNs (*magenta*) are visualized in 3D in overlap with the original Oprm1-Cre:tdTomato signal (*white*). Dynamic scale bar in the bottom left corner.

